# Aurora kinase Ipl1 facilitates bilobed distribution of clustered kinetochores to ensure error-free chromosome segregation in *Candida albicans*

**DOI:** 10.1101/542605

**Authors:** Neha Varshney, Kaustuv Sanyal

**Affiliations:** Molecular Mycology Laboratory, Molecular Biology & Genetics Unit, Jawaharlal Nehru Centre for Advanced Scientific Research, Jakkur, Bangalore 560064, India

**Keywords:** Microtubules, kinetochore geometry, ploidy, anti-fungal drug resistance, kinetochore-microtubule attachments

## Abstract

*Candida albicans,* an ascomycete, has an ability to switch to diverse morphological forms. While *C. albicans* is predominatly diploid, it can tolerate aneuploidy as a survival strategy under stress. Aurora kinase B homolog Ipl1 is a critical ploidy regulator that controls microtubule dynamics and chromosome segregation in *Saccharomyces cerevisiae*. In this study, we show that Ipl1 in *C. albicans* has a longer activation loop than that of the well-studied ascomycete *S. cerevisiae*. Ipl1 localizes to the kinetochores during the G1/S phase and associates with the spindle during mitosis. Ipl1 regulates cell morphogenesis and is required for cell viability. Ipl1 monitors microtubule dynamics which is mediated by separation of spindle pole bodies. While Ipl1 is dispensable for maintaining structural integrity and clustering of kinetochores in *C. albicans*, it is required for the maintenance of kinetochore geometry to form bilobed structures along the mitotic spindle, a feature of Ipl1 that was not observed in other yeasts. Depletion of Ipl1 results in erroneous kinetochore-microtubule attachments leading to aneuploidy-associated resistance to fluconazole, the most common anti-fungal drug used to treat *Candida* infections. Taking together, we suggest that Ipl1 spatiotemporally ensures kinetochore geometry to facilitate bipolar spindle assembly crucial for ploidy maintenance in *C. albicans*.

## Introduction

Faithful inheritance of the duplicated genetic material to the daughter cells during cell division relies on the process of accurate chromosome segregation. Members of the Aurora family of serine/threonine protein kinases regulate various processes during chromosome segregation such as chromosome condensation, kinetochore-microtubule (MT) interactions, spindle-assembly checkpoint (SAC), spindle dynamics and cytokinesis (Andrews. *et al.*, 2003, Carmena. & Earnshaw., 2003). In ascomyceteous budding yeast *Saccharomyces cerevisiae*, Ipl1 is the only Aurora protein kinase which confers the **I**ncrease-in-**pl**oidy phenotype (Francisco & Chan, 1994, Chan & Botstein, 1993, Biggins. *et al.*, 1999). The protein comprises of a highly conserved C-terminal catalytic domain and a diverged N-terminal domain. The C-terminal catalytic domain of the protein contains an activation loop motif between subdomains VII and VIII, and a destruction-box to direct the Anaphase Promoting Complex (APC)-dependent degradation (Giet & Prigent, 1999). Ipl1, as a part of Chromosome Passenger Complex (CPC), associates with Sli15, Bir1 and Nbl1, orthologues of the human Inner centromere protein INCENP, Survivin and Borealin respectively, and exhibits a dynamic localization throughout the cell cycle (Buvelot. *et al.*, 2003).

During chromosome segregation, the sister chromatids must attach to the opposite spindle poles to achieve bi-orientation before the onset of anaphase to segregate appropriately in daughter cells. Ipl1 senses the tension across the centromere (*CEN*), stabilizes the bi-oriented kinetochore-MT attachments and destabilizes the erroneous kinetochore-MT attachments by regulating the phosphorylation of kinetochore proteins such as Dam1, Ndc80 etc. (Kim. *et al.*, 1999, Tanaka. *et al.*, 2002, Cheeseman. *et al.*, 2002). In addition, Ipl1 has been shown to associate with the mitotic spindle to regulate its timely disassembly by phosphorylating microtubule-associated proteins (MAPs) such as Bim1 and Ase1 (Kang. *et al.*, 2001, Zimniak. *et al.*, 2009, Woodruff. *et al.*, 2010).

*Candida albicans,* is the most prevalent human fungal pathogen, accounting for approximately 400,000 life-threatening infections per year (Brown *et al.*, 2012). While it is a human commensal, it becomes an opportunistic pathogen and resides in the host with reduced immune competence or an imbalance of competing bacterial microflora (Berman, 2012). Azole-associated acquisition of aneuploidy is a well-elucidated mechanism that provides fitness during anti-fungal drug resistance in *C. albicans* (Selmecki *et al.*, 2009). One of these aneuploid states is the isochromosome 5L (i(5L)) which has been shown to confer fluconazole resistance due to amplification of genes such as *TAC1* and *ERG11* present on the left arm of chromosome 7 (Selmecki *et al.*, 2006, Selmecki *et al.*, 2008). The virulence of the organism has been shown to be associated with the morphological transition between yeast and hyphal cells. In these morphological forms, the MT-associated motor proteins have been shown to regulate nuclear movements, chromosome segregation, and cytoskeleton remodeling (Finley *et al.*, 2008, Martin *et al.*, 2004, Sherwood & Bennett, 2008, Frazer *et al.*, 2015). Yeast and filamentous hypha, both exhibit variations in the length of the mitotic spindle, the former having shorter mitotic spindles of maximum of 8 µm and the latter with a longer spindle MTs of maximum of 20 µm, indicating that the cell morphology influences the length of MTs in the same organism (Barton & Gull, 1988, McCoy *et al.*, 2015). *C. albicans* genome encodes only one form of the kinesin-5 (Kip1) and kinesin-14 (Kar3Cik1) family motors in contrast to the two kinesin-5 (Kip1 and Cin8) and kinesin-14 (Kar3Cik1 and Kar3Vik1) family motors in *S. cerevisiae* (Chua *et al.*, 2007, Frazer *et al.*, 2015). *C. albicans* exhibits various cellular morphologies and provides an ideal system to explore various features of MT dynamics during mitosis because of its extraordinary genomic plasticity to tolerate chromosomal alterations and ploidy changes under unfavorable conditions such as the presence of anti-fungal drugs (Sanyal, 2012).

The primary events of the mitotic cell cycle are largely conserved between *S. cerevisiae* and *C. albicans* both of which belong to the same phylum of Ascomycota although these two-yeast species diverged from a common ancestor >600 million years ago. The SPBs are embedded in the nuclear envelope (NE) that never breaks down resulting in a closed mitosis. The kinetochores remain clustered and are bound to the MTs throughout the cell cycle (Sanyal & Carbon, 2002, Roy *et al.*, 2011). Further, the correlation of “one MT/kinetochore” established in *S. cerevisiae* having 125 bp short point *CENs* seems to be conserved in *C. albicans* having 3-5 kb long CENP-A rich *CENs* (Joglekar *et al.*, 2008). However, in contrast to *S. cerevisiae,* the kinetochore architecture in *C. albicans* is stabilized in a coordinated interdependent manner by its individual components indicating that the kinetochore may not be a layered structure in *C. albicans* (Thakur & Sanyal, 2012). Depletion of essential kinetochore proteins from the different layers of the kinetochore results in kinetochore unclustering followed by a complete collapse of the kinetochore architecture and reduction in cellular levels of the centromere-specific histone CENP-A (Thakur & Sanyal, 2012, Roy *et al.*, 2011).

Here, we explored the function of Aurora kinase Ipl1 in the spatiotemporal regulation of the MT dynamics during chromosome segregation in *C. albicans*. Using a conditional promoter, we modulated the cellular levels of Aurora B kinase Ipl1 and demonstrated that Ipl1 regulates MT dynamics. Ipl1 maintains bilobed kinetochore organization and its geometry to ensure proper separation of chromatin to prevent aneuploidy-associated drug resistance.

## Results

### A homolog of Ipl1 in C. albicans has a distinct activation loop

C6_02320C, annotated as the homolog of Ipl1 in *C. albicans* (CaIpl1) in the Candida Genome Database (http://www.candidagenome.org/), was identified through a BLAST search using the primary amino acid sequence of Ipl1 in *S. cerevisiae* as the query. The sequence of putative CaIpl1 encodes a serine-threonine protein kinase of 530 amino acids long peptide with a predicted molecular weight of 62 kDa that shares a high degree of amino acid sequence similarity to Aurora kinases from other organisms (Supporting Information, Fig. S1) (Robert & Gouet, 2014). Aurora kinases have a characteristic catalytic domain which contains a conserved motif (DFGWSXXXXXXX-RXTXCGTXDYLPPE) in the activation loop domain between subdomains VII and VIII and a D2-type destruction box (LLXXXPXXRXXLXXXXXHPW), near its C terminus (Giet & Prigent, 1999). Strikingly, CaIpl1 contains an extended activation loop domain (residues 384-434) within the conserved catalytic domain (Supporting Information, Fig. S1A). In addition, it also contains the conserved destruction box sequence (residues 496-515) as a part of catalytic domain. We constructed a phylogenetic tree with putative Ipl1 protein sequences from the members the *Saccharomycotina* species complex (Supporting Information, Fig. S1B). *C. albicans* belongs to the CTG clade in which the CTG codon is often translated as serine, rather than leucine (Turner & Butler, 2014). CaIpl1 kinase branches off near the base of the clade consisting of the family *Pichiaceae*. Thus, our analysis suggests that Ipl1 protein in the CTG clade species significantly diverged from the same in the well-studied yeast *S. cerevisiae* (ScIpl1).

### Dynamic localization of Ipl1 during different stages of cell cycle in C. albicans

To determine whether Ip11 functions differently in *C. albicans* from that of the *S. cerevisiae*, we examined the localization of Ipl1 in *C. albicans*. We functionally expressed Ipl1-Protein-A from its native promoter in the strain *IPL1-TAP-URA3/IPL1* (CNV1) (Puig *et al.*, 2001). Ipl1-Protein-A could be detected by western blot analysis using lysates obtained after immunoprecipitation with IgG sepharose beads (Supporting Information, Fig. S2). However, Ipl1 was found undetected by western blot analysis using lysates obtained prior to immunoprecipitation, indicating that Ipl1 is expressed at a low level in *C. albicans*. Further, to visualize its subcellular localization, we functionally expressed it as a fusion protein with 2xGFP at its C-terminus under the native promoter of *IPL1* in the strain *IPL1p-IPL1-2xGFP-HIS1/ipl1::FRT, NDC80/NDC80-RFP-ARG4* (CNV3) where the kinetochore protein Ndc80 was tagged with RFP and co-expressed. Ipl1 co-localizes to the kinetochores during the S-phase and metaphase (Fig. 1A). However, Ipl1 decorates the mitotic spindle in anaphase but no nuclear localization of Ipl1 could be detected in post-anaphase. In unbudded cells, Ipl1 was occasionally detected as a dot-like structure. Thus, Ipl1 is localized to the kinetochore in a cell-cycle-dependent manner in *C. albicans,* which is distinct from Ipl1’s localization in *S. cerevisiae* (Buvelot. *et al.*, 2003, Tanaka. *et al.*, 2002, Biggins. *et al.*, 1999). We also studied the localization of Ipl1-2xGFP in nocodazole-treated metaphase-arrested CNV3 cells. Majority of these cells exhibited kinetochore localization of Ipl1 indicating that Ipl1 localizes to the kinetochore under tension in *C. albicans* (Fig. 1B). We also noticed a higher accumulation of Ipl1 at the kinetochore when kinetochore-MT interaction is perturbed in nocodazole-treated cells (Fig. 1C).

**Figure 1.**
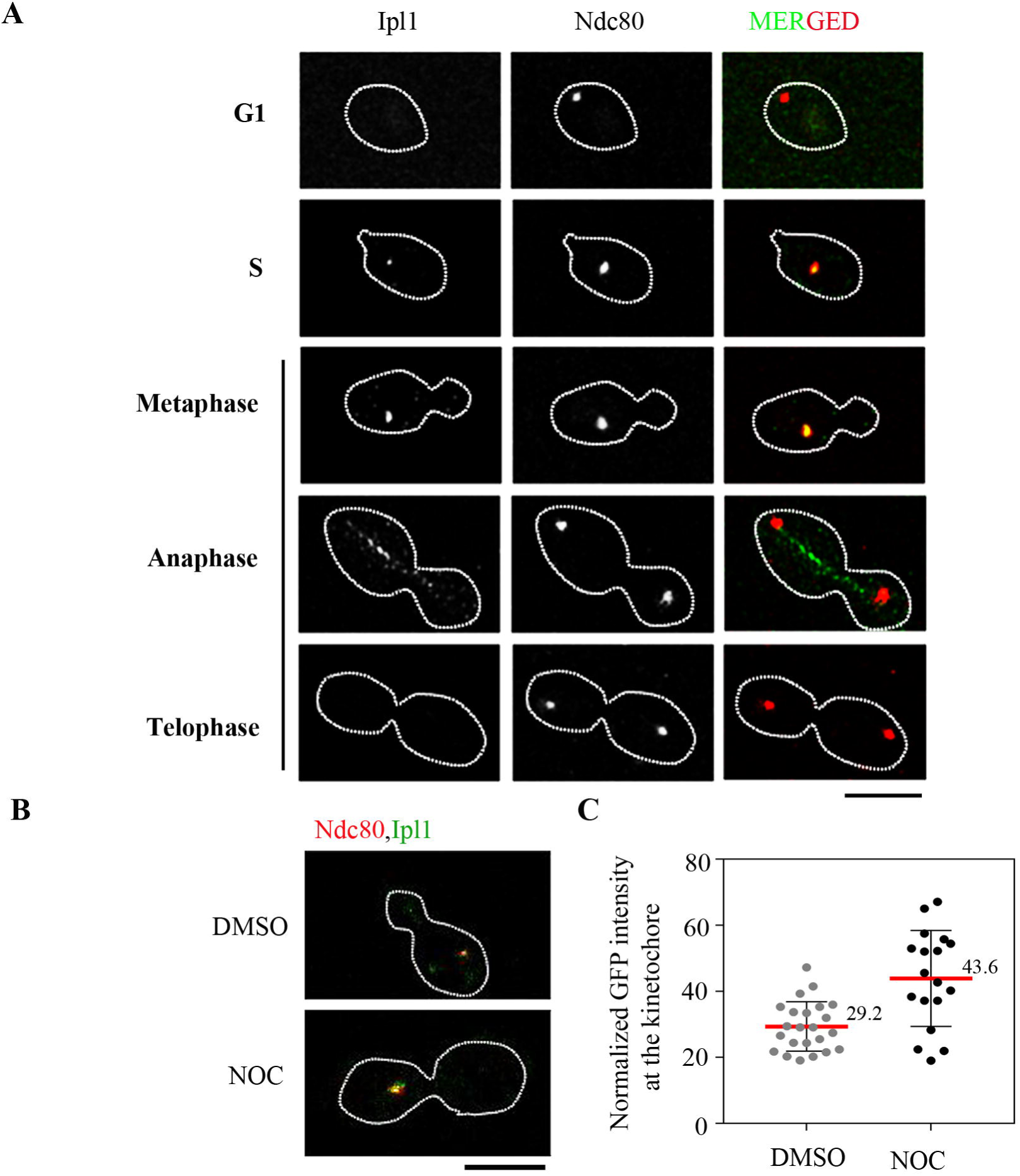
Dynamic localization of Ipl1 in *C. albicans*. (**A)** Localization of Ipl1 in the wild-type CNV3 cells co-expressing Ipl1-2xGFP and a kinetochore marker, Ndc80-RFP in interphase and mitotic cells. During S-phase and metaphase, Ipl1 co-localizes with Ndc80. During anaphase, Ipl1 localizes to the spindle. Bar, 5μm. **(B)** Localization of Ipl1 in the wild-type CNV3 cells co-expressing Ipl1-2xGFP and a kinetochore marker, Ndc80-RFP upon treatment with DMSO and nocodazole (NOC), 20 µg/ml. **(C)** Quantification of Ipl1-2xGFP signal intensities at the kinetochores in the DMSO and NOC treated wild-type cells.

### Depletion of Ipl1 leads to a loss in viability and multiple morphological defects

To study the function of Ipl1 in *C. albicans*, we constructed its conditional mutant strain by placing *IPL1* under the regulatable *PCK1* promoter which is repressed in the presence of glucose (non-permissive) but expressed in the presence of succinate (permissive) (Fig. 2A) (Leuker *et al.*, 1997). Two independent conditional *ipl1* mutants *PCK1p-IPL1/ipl1::FRT* (CNV6 and CNV7) were constructed and were confirmed by Southern blot analysis (Supporting Information, Fig. S3A). The wild-type *IPL1*/*IPL1* (RM1000AH) and the conditional *ipl1* mutants, *PCK1p-IPL1/ipl1::FRT* (CNV6 and CNV7), when streaked for single colonies on the plates revealed that cells having depleted levels of Ipl1 could not form colonies with the same efficiency as that of the wild-type (Fig. 2A). Ipl1-depletion cells consistently produced smaller colonies of variable sizes as compared to those of wild-type strain (Fig. 2B). Next, we examined whether Ipl1 plays a role in regulating morphogenesis in *C. albicans* by growing cells of *PCK1p-IPL1/ipl1::FRT* (CNV6) under non-permissive conditions. Microscopic observations revealed that the majority of Ipl1-depleted cells were larger in size than the wild-type cells with altered cellular morphologies (Fig. 2C). We quantified different morphological phenotypes obtained upon depletion of Ipl1 and classified them as dividing (unbudded, small-budded, large-budded and elongated), terminal, branched, chained and complex (cells with two or more branches and trimeras) cells. Importantly, in contrast to the wild-type cells, the Ipl1 depletion resulted in the significant accumulation of enlarged, branched, terminal and complex multimeric cells after 8 h of growth under non-permissive conditions (Fig. 2D), suggesting that Ipl1 depletion triggers altered morphogenesis in *C. albicans.* A viability assay was performed to determine if the altered morphologies contribute to increased cell death in Ipl1-depleted cells. A significant reduction (~70%) in the viability of Ipl1-depleted cells after 8 h of growth in non-permissive conditions as compared to the wild-type cells revealed that Ipl1 is required for viability in *C. albicans* (Supporting Information, S3B). This result also explains our inability to obtain a null mutant of *ipl1*. Since Ipl1-depleted cells exhibited a significant drop in viability associated with dramatic morphological changes after 8 h of growth under non-permissive conditions, we performed subsequent experiments under similar conditions (unless specified) to study functions of the mutant *ipl1*.

**Figure 2.**
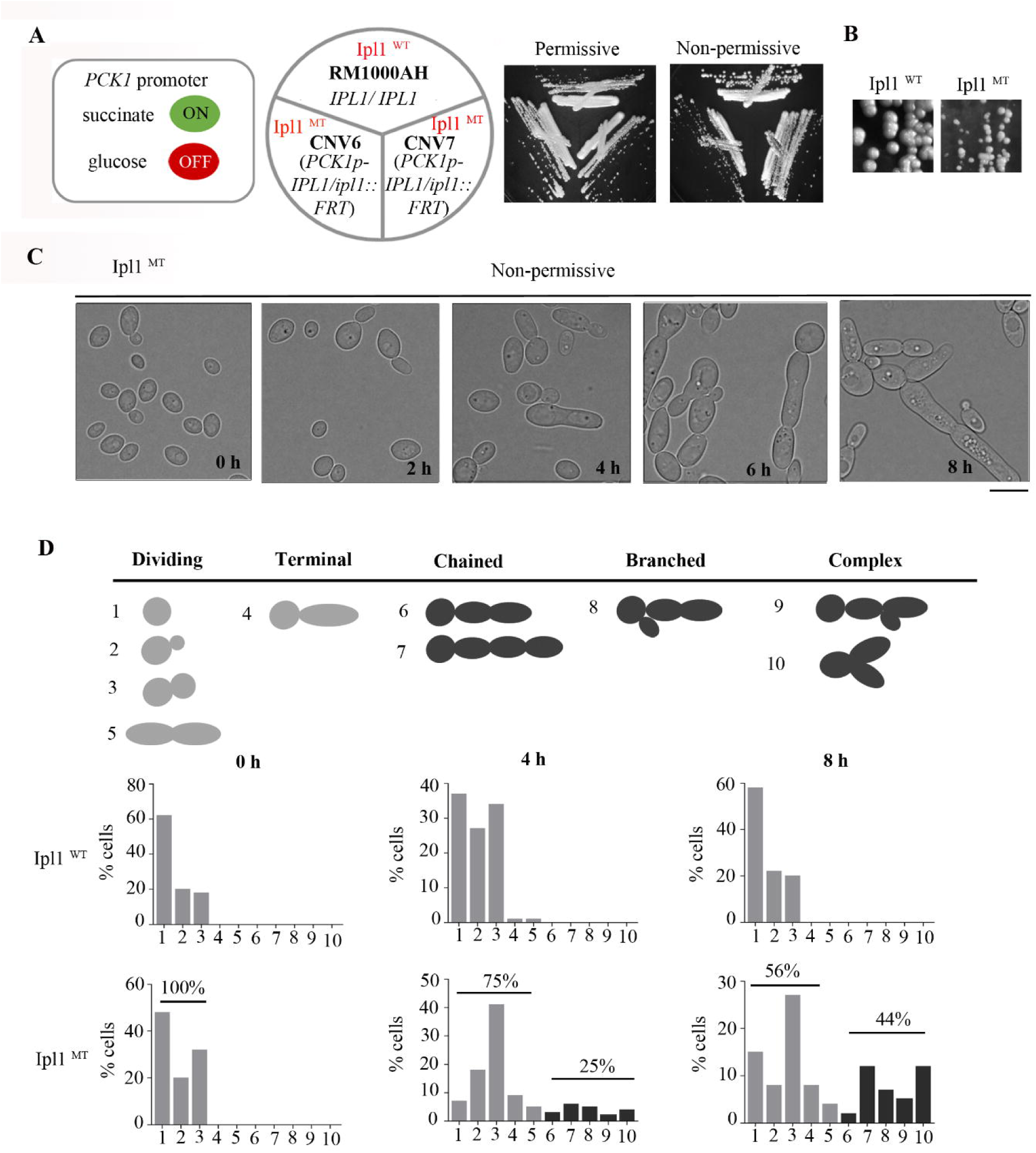
Ipl1 depletion induces aberrant morphological states in *C. albicans*. **(A)** The regulation of the *PCK1* promoter used to control the expression of *IPL1*. The cells of wild-type RM1000AH and Ipl1 depletion mutants, CNV6 and CNV7 were streaked on the plates containing permissive and non-permissive media. The plates were photographed after 3 days of incubation at 30°C. **(B)** Reduction in the size of colonies in Ipl1-depleted CNV6 cells as compared to the wild-type RM1000AH cells in the non-permissive conditions. **(C)** Microscopic images of the Ipl1 mutant cells of CNV6 after growth in non-permissive conditions for the indicated time-points. Scale 5µm. **(D)** The diagrammatic representation and quantification of enlarged and complex morphological phenotypes obtained in Ipl1-depleted cells of CNV6 as compared to the wild-type cells of RM1000AH after growth in the non-permissive conditions at the indicated time intervals.

### Depletion of Ipl1 induces the assembly of aberrant spindles with inappropriately separated spindle poles

Next, we set out to determine why Ipl1-depleted cells exhibit severely altered morphologies of the organism. The controlled disassembly and reassembly of MT fibers regulate the architecture or the shape of a cell. It has been shown previously MT motor mutants of *kip1*, *kar3,* or *cik1* result in altered cellular morphologies in *C. albicans* (Frazer *et al.*, 2015, Chua *et al.*, 2007, Sherwood & Bennett, 2008). To test whether the aberrant morphologies in the Ipl1-depleted cells are due to altered MT dynamics, we first depleted Ipl1 in two independent mutant strains, *ipl1::FRT/PCK1*p*-IPL1* (CNV6 and CNV7) for 4 h and spotted them on the plates containing thiabendazole (40 µg/ml), a MT-depolymerizing agent. It must be noted that the depletion of Ipl1 for this experiment was done for only 4 h to avoid the formation of cells with altered cellular morphologies that could have affected the proper estimation of the cell number. Retardation in the growth of the Ipl1-depleted cells as compared to the wild-type in the presence of thiabendazole indicates that depletion of Ipl1 affects MT dynamics (Fig. 3A). Shifts in the temperatures are also known to modulate the depolymerization and polymerization dynamics of the MTs.

**Figure 3.**
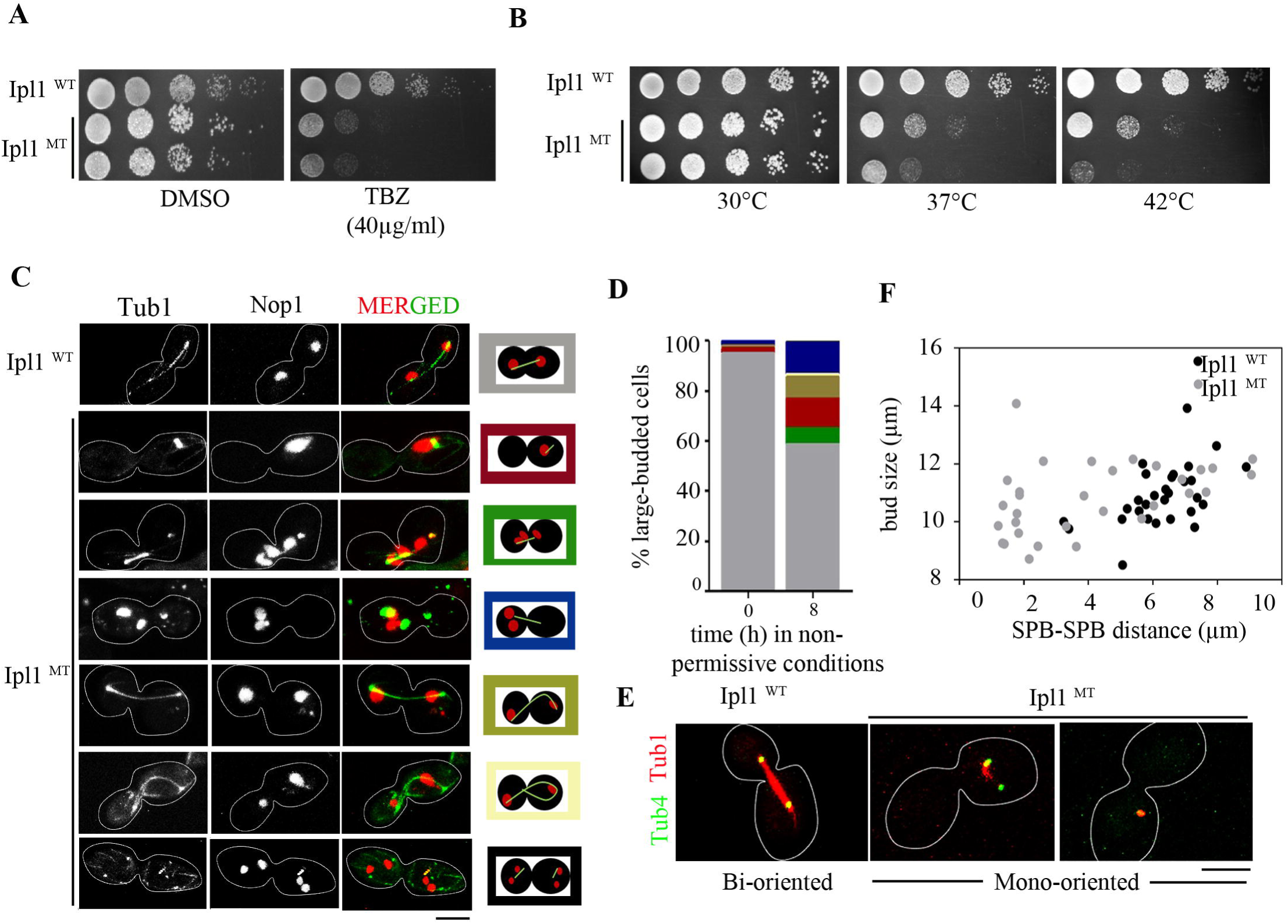
Ipl1-depleted cells are defective in the microtubule dynamics and spindle pole body (SPB) separation. **(A)** The thiabendazole (TBZ)-sensitivity assay depicting the altered dynamics of the MTs in Ipl1-depleted cells grown in the non-permissive conditions for 4 h. The wild-type RM1000AH and Ipl1-depleted CNV6 and CNV7 cells were 10-fold serially diluted and spotted on the non-permissive medium containing TBZ (40µg/ml). DMF was used as a no-drug control. (**B)** The temperature-sensitivity assay corroborating the perturbed MT structures in Ipl1-depleted cells grown in the non-permissive conditions for 4 h. (**C)** Microscopic images of cells having GFP-Tub1 and RFP-Nop1 depicting structural changes of MTs and nucleolus in the Ipl1-depleted CNV9 cells as compared to wild-type 12856. Bar, 5µm. Cartoons representing different types of MTs with respect to the nucleolus observed in the wild-type and mutant cells (right). (**D)** Quantification of MTs of altered structures in the conditional mutant of *ipl1* after growth in the non-permissive conditions for 8 h. (**E**) Microscopic images of cells having GFP-Tub4 and RFP-Tub1 depicting the distance between the two SPBs in the pre-anaphase cells of the wild-type strain CNV13 and Ipl1-depletion mutant strain CNV14. Chromosomes in majority of the wild-type cells exhibit bipolar attachments while significant proportion of the Ipl1-depleted cells exhibit chromosomes with that lack biploar attachments. Bar, 5µm. (**F)** Quantification of the distances between the SPBs measured in the wild-type and Ipl1-depleted pre-anaphase cells having the bud-size of 8-16 µm.

To confirm whether the MT dynamics is perturbed in the Ipl1-depleted cells, we spotted the Ipl1-depleted cells (CNV6 and CNV7) and the wild-type cells (RM1000AH) on glucose plates and incubated them at variable temperatures (30°C-42°C). As compared to the optimal temperature (30°C), Ipl1-depleted cells exhibited retarded growth at the higher temperatures (37°C and 42°C) (Fig. 3B). This result was similar to the growth defect observed in the *ipl1-2* mutant in *S. cerevisiae* and *kar3Δ/kar3Δ* mutant in *C. albicans* cells at higher temperatures (Sherwood & Bennett, 2008, Robinson *et al.*, 2012). However, mutant of *ipl1* in *S. cerevisiae, ipl1-315* having reduced kinase activity displayed a different phenotype as these mutant cells could grow at 37°C (Kotwaliwale. *et al.*, 2007).

Next, we sought to determine the possible reasons for exhibiting severe defects in the MT dynamics by Ipl1-depleted cells in *C. albicans*. To test this, we first depleted Ipl1 for 8 h in cells of *TUB1-GFP URA3/TUB1, NOP1-RFP NAT/NOP1, PCK1p-IPL1-HIS1/ipl1::ARG4* (CNV9) that expresses α-tubulin Tub1 tagged with GFP and nucleolar marker Nop1 tagged with RFP, and analyzed the morphology of the MTs. We observed that as compared to the wild-type strain *TUB1-GFP URA3/TUB1 NOP1-RFP NAT/NOP1, IPL1/IPL1* (12865), both the morphology and dynamics of MTs were significantly affected in the *ipl1* mutant cells (Fig. 3C). While close to 50% Ipl1-depleted cells exhibited apparently normal MTs, a fraction of population exhibited an array of spindle defects ranging from a) short spindle with unsegregated chromatin (15%), b) short spindle with mis-segregated chromatin (6%), c) aberrant mitotic spindle with mis-segregated chromatin in the mother cell (17%), d) long spindle (8%), and, e) forked-shaped or broken spindle (2%) (Fig. 3D). Since most of the defective spindles were short, we scored the mitotic spindle length by measuring SPB to SPB distance in the pre-anaphase cells of the wild-type strain *TUB4-GFP-URA3/TUB4, TUB1-RFP-HIS1/TUB1, IPL1/IPL1* (CNV13) and Ipl1-depleted strain *TUB4-GFP-URA3/TUB4 TUB1-RFP-HIS1/TUB1 ipl1::FRT/PCK1*p*-IPL1* (CNV14) co-expressing GFP-tagged Tub4 and RFP-tagged Tub1 (Fig. 3E). The cells having the bud-size (length of the major axis of the cell) of 10-12 µm are considered as the pre-anaphase cells. Remarkably, while the wild-type pre-anaphase cells had SPBs well-separated from each other (6-8 µm), Ipl1-depleted pre-anaphase cells had SPBs localized significantly closer to each other (2-4 µm) (Fig. 3F). However, we also observed single or two very closely spaced foci of SPBs (Tub4-GFP), co-localizing with the spindle (Tub1-RFP) indicating that Ipl1 depletion possibly results in the formation of mono-oriented spindles in a smaller percentage of cells. Together, these results indicate that the Ipl1-depleted cells have difficulties in separating SPBs that results in a mitotic delay or cell death.

### Ipl1 is involved in the bilobed organization of the kinetochores and their attachment with the microtubules

MTs with an opposite polarity overlap in a zone between the two SPBs and are pushed apart by specialized motors such as plus-end directed motors of the kinesin-5 family, Kip1, and Cin8. Kip1 and Cin8 work together to cluster kinetochores in *S. cerevisiae* (Tytell & Sorger, 2006). In addition, kinesin-8 family, Kip3 is required for kinetochore positioning along the metaphase spindle (Wargacki *et al.*, 2010). Thus, we were curious to test whether the organization or spatial geometry of kinetochores is affected in the Ipl1-depleted cells having aberrantly spaced SPBs. To investigate the defects in kinetochore geometry and arrangement, we first depleted Ipl1 for 8 h in cells co-expressing GFP-tagged Dad2 (outer kinetochore) and RFP-tagged Tub1, *DAD2-GFP-URA3/DAD2, TUB1-RFP-HIS1/TUB1, ipl1::FRT/PCK1*p-*IPL1* (CNV19), analysed kinetochore (Dad2-GFP) signals along the spindle-axis (Tub1-RFP), and compared them with the wild-type cells *DAD2-GFP-URA3/DAD2, TUB1-RFP-HIS1/TUB1, IPL1/IPL1* (CNV22). We observed that Ipl1-depleted cells contained mis-positioned Dad2-GFP foci along the spindle-axis or had abnormally diffuse GFP lobes as compared to the bilobed distribution of the Dad2-GFP foci observed in the wild-type cells along the spindle-axis (Fig. 4A). The peak fluorescence intensity of the Dad2-GFP signals was towards the poles in the wild-type, while the peak fluorescence intensity of the Dad2-GFP was shifted towards the spindle equator or along the length of the spindle in the large-budded Ipl1-depleted cells (Fig. 4B). In addition, we also observed unequally separated kinetochore clusters in minor percentage of Ipl1-depleted cells. Unequally separated kinetochore clusters at the mother-daughter cell junction could be due to an accumulation of MTs with unrectified mono-oriented attachments generated in the cell. Further, we estimated the number of large-budded Ipl1-depleted cells with defective kinetochore geometry and kinetochore-microtubule attachments. We found that close to 50% Ipl1-depleted cells were defective in distribution of the kinetochore clusters (Supporting Information, Fig. S3A and S3B). We have shown previously that the partial depletion of essential kinetochore proteins affects the integrity of the kinetochore cluster and stability of the centromeric histone CENP-A in *C. albicans* (Thakur. & Sanyal., 2012). Therefore, we further tested whether the mis-organization of kinetochore clusters obtained upon depletion of Ipl1 also affects the stability and integrity of the centromeric histone CENP-A. We prepared lysates from *CENP-A/CENP-A-TAP-URA3, ipl1::FRT/PCK1p-IPL1-NAT* cells (CNV17) grown in non-permissive conditions for various time intervals, and performed western blot analysis to measure the levels of protein A-tagged CENP-A. We find that the total cellular protein levels of protein A-tagged CENP-A remain unaltered upon depletion of Ipl1 (Fig. 4C), indicating that the stability of CENP-A remains unaffected upon depletion of Ipl1. Further, we observed no significant difference in the total mean intensity of CENP-A-GFP signals per kinetochore cluster in the Ipl1-depleted cells *CENP-A/CENP-A:GFP:CENP-A, ipl1::FRT/PCK1p-IPL1* of CNV27 as compared to the wild-type cells *CENP-A/CENP-A:GFP:CENP-A* (YJB8675) indicating that the kinetochore localization of CENP-A remains unaltered upon Ipl1 depletion. Similarly, we tested the difference in the occupancy of CENP-A at the *CENs* in the presence and absence of Ipl1 by performing chromatin immunoprecipitation (ChIP) in the strains CAKS102 and CNV17. We find that the occupancy of CENP-A at the *CEN*s remains unaffected in the Ipl1-depleted cells (Fig. 4D and 4E). Together, these lines of evidence indicate that Ipl1 is required for maintaining the kinetochore geometry during the metaphase-anaphase transition but dispensable for the integrity of kinetochores in *C. albicans*.

**Figure 4.**
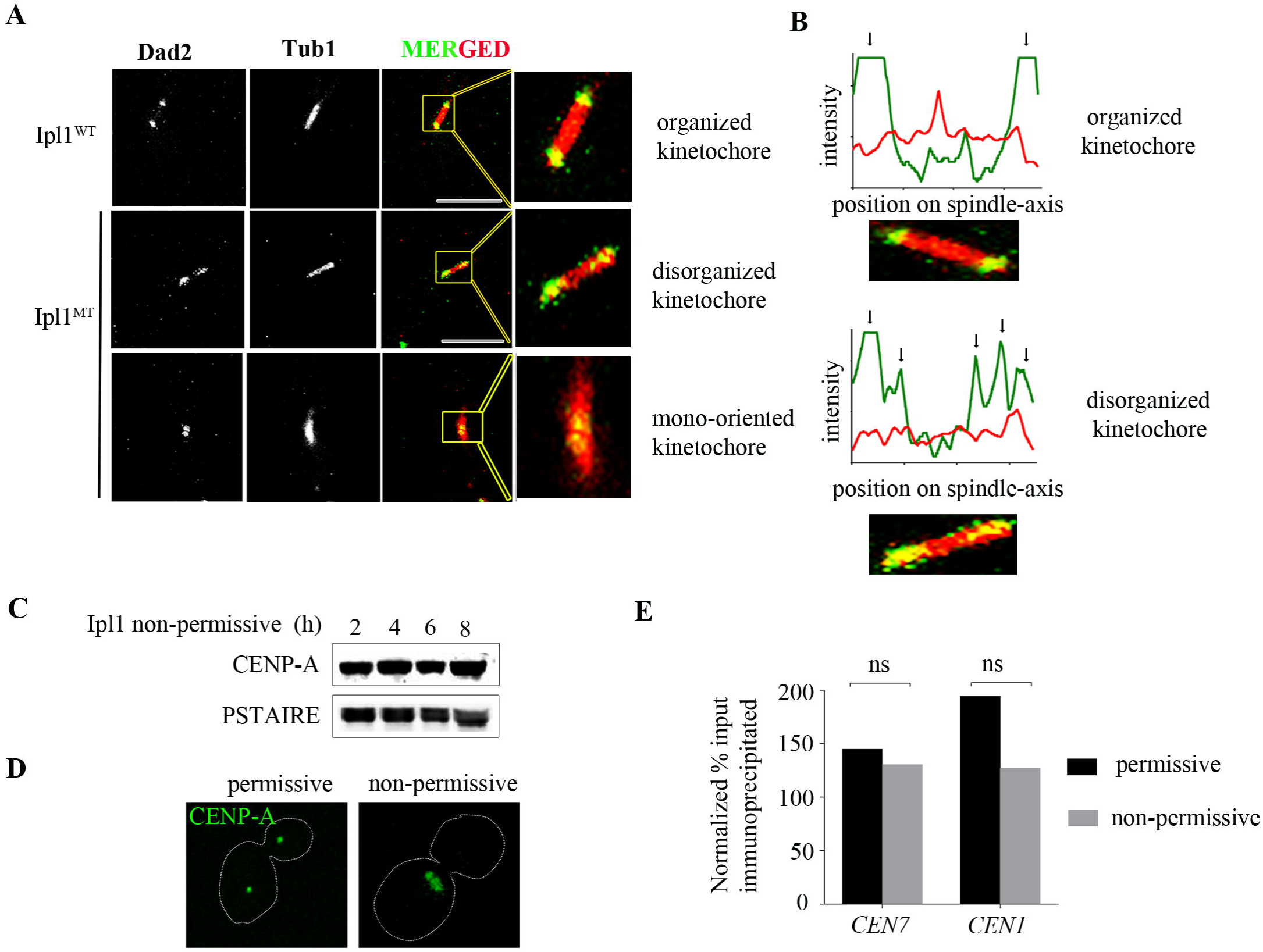
Depletion of Ipl1 leads to disorganization and improper attachment of kinetochores. **(A)** Images of the organization of kinetochores (Dad2-GFP) along the spindle axis (Tub1-RFP) in the wild-type CNV21 and *ipl1* mutant CNV22 cells. The kinetochores remained clustered in the large-budded wild-type cells with the peak fluorescence intensity towards the spindle poles, while the kinetochores were declustered/disorganized with the peak fluorescence intensity shifted towards the spindle mid-zone in the large-budded Ipl1-depleted cells (zoomed). (**B)** Histogram plot indicated the fluorescence intensity of Tub1-RFP (red) and Dad2-GFP (green), (indicated with black arrows) along the spindle axis in the wild-type and Ipl1 mutant cells showing organized and disorganized kinetochores respectively. (**C)** The levels of CENP-A expression in cell lysates prepared from CNV17 cells after the indicated time of incubation in non-permissive media conditions. Western blot analysis was done using anti-TAP and anti-PSTAIRE antibodies. **(D)** Images of CENP-A-GFP localization in CNV27 cells grown in permissive and non-permissive conditions. **(E)** Quantitative real-time PCR (qPCR) were performed in the *ipl1* mutant strain grown in permissive and non-permissive conditions for 8 h for enrichment of CENP-A-Prot-A at the core *CEN7* and *CEN1* after normalizing with the non-*CEN*. Enrichment of CENP-A at the *CEN* was calculated as a percentage of the total chromatin input and values were plotted as mean of two independent experiments (three technical replicates for each experiment).

### Ipl1 depletion results in nuclear mis-segregation which results in aneuploidy-associated drug resistance

Defects in the kinetochore-MT attachments could result in the altered distribution of kinetochores in Ipl1-depleted cells. To test whether defective kinetochore-MT attachments result in altered ploidy in *C. albicans,* we sought to visualize the segregation of the duplicated chromatin mass by analyzing the fluorescence of H2B-tagged with GFP in the wild-type cells, *HTB*/*HTB*-*GFP*::*SAT1 IPL1/IPL1* (RSY15) and Ipl1-depleted strain *HTB*/*HTB*-*GFP*::*SAT1 ipl1::ARG4/PCK1p-IPL1-HIS1* (CNV24) after growing them in the non-permissive conditions for 8 h. Depletion of Ipl1 severely affected the dynamics of nuclear segregation in both budded and elongated yeast cells (Fig. 5A and 5B). Ipl1-depleted cells exhibited five distinct nuclear segregation phenotypes in a similar ratio a) unsegregated chromatin in the small-budded cells, b) segregated chromatin between mother and daughter cell, c) unsegregated chromatin in the large-budded cells with chromatin present in either the mother cell or unequally segregated chromatin in mother and daughter bud, d) stretched chromatin in large-budded cells, e) aberrantly segregated chromatin in pseudo-hyphal/elongated/multi-budded cells. In contrast, the majority of wild-type cells exhibited either unsegregated chromatin before division in the small-budded cell or equal segregation of the chromatin mass between mother and daughter cell. A dramatic increase in the number of large-budded cells having unsegregated or stretched chromatin in the mutant suggested that there could be a delay in the segregation of chromatin as a stretched chromatin mass persists for a longer duration in the mutant cells. To understand the occurrence of these phenotypes, we monitored the dynamics of separation of the chromatin mass in the live wild-type cells of *HTB*/*HTB*-*GFP*::*SAT1 IPL1/IPL1* (RSY15) and mutant Ipl1-depleted strain, CNV24 cells by live-cell imaging for 70 min from the time of inception of the daughter bud (Fig. 5C, Supporting Information, Movie S1). We observed that Ipl1-depleted cells exhibit unsegregated or stretched chromatin at the mother-daughter cell junction that persists for a longer duration as compared to the wild-type. However, there was no significant difference observed between the average time required for the segregation of chromatin from the time of the inception of bud in the wild-type cells (mean = 48.5 min) as compared to the subset Ipl1-depleted cells (mean = 50.3 min) which could finally manage to divide the chromatin mass (Fig. 5D). Despite extended time spent as a stretched chromatin, how some of the Ipl1-depleted cells segregate the chromatin mass within the same time frame as that of the wild-type cells is intriguing and requires further analysis. In conclusion, a delay in the separation of chromatin or prolonged stretching of chromatin could possibly result in unequal segregation of the nucleus resulting in death or aneuploidy in Ipl1-depleted cells.

**Figure 5.**
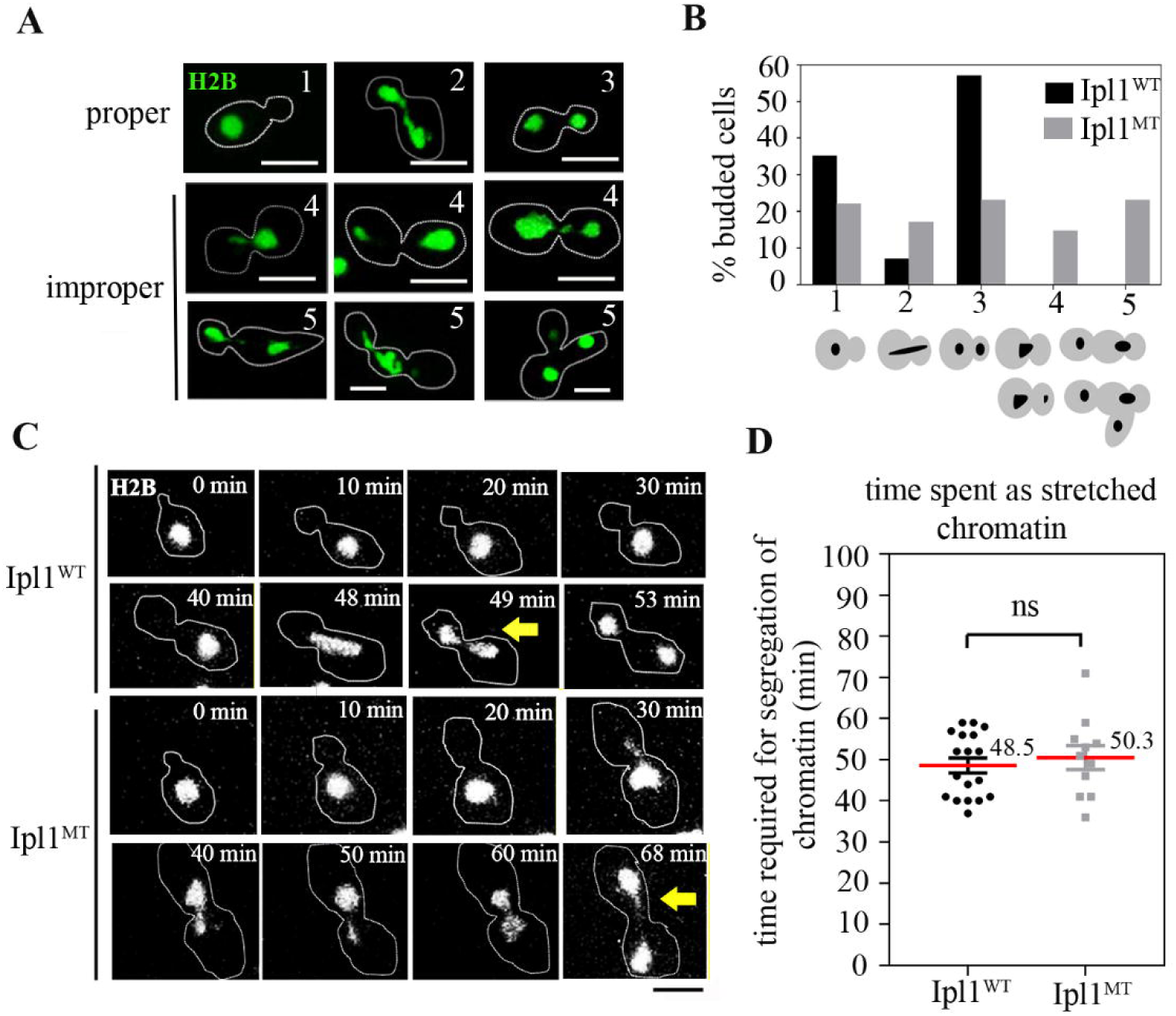
Defective separation of chromatin in Ipl1-depleted cells. **(A)** Microscopic analysis of the distribution of chromatin between mother and daughter buds in the budded cells of wild-type RSY15 and Ipl1-depletion mutant CNV24 carrying histone H2B-GFP. All the phenotypes were classified into five classes (1-5) Bar 5µm. **(B)** Quantification of the chromatin distribution phenotypes in the wild-type and Ipl1-depleted cells classified into five classes (1-5, shown below the bar graph). Cartoons below each class show cellular morphologies (gray) the chromatin mass (black). **(C)** Snapshots from a time-lapse movie of nuclear segregation in the wild-type and Ipl1-depleted cells at the indicated time point from the time of inception of bud till the segregation. The nucleus is marked by histone H2B-GFP. Bar 5µm. The formation of the stretched chromatin mass is marked with a yellow arrow in each case. **(D)** Quantification of the time (min) required for chromatin segregation during cell division from the inception of bud in the wild-type and Ipl1-depleted cells.

*C. albicans* exhibits aneuploidy as an adaptive mechanism in response to stress such as fluconazole (FLC), the most common drug used for treating *Candida* infections. Therefore, we tested whether Ipl1-depleted cells carrying aneuploidy display FLC resistance. For this, we estimated the presence of aneuploidy in Ipl1-depleted cells grown in non-permissive conditions by flow cytometric analysis (FACS). The existence of aneuploid cells (>4n) in a population of Ipl1-depleted cells in the strain CNV6 within 4 h of protein depletion further confirmed that Ipl1 is required for ploidy maintenance in *C. albicans* (Fig. 6A).. Further, to test whether Ipl1-depleted cells carrying aneuploidy are resistant to FLC, we depleted Ipl1 in two independent mutants, CNV6 and CNV7 for 4 h and spotted them on the plates containing FLC (64 µg/ml) along with the wild-type strain, RM1000AH. Indeed, Ipl1-depleted cells displayed a higher number of FLC resistant colonies as compared to the wild-type (Fig. 6B). Next, we quantified aneuploidy by performing a chromosome loss assay in Ipl1-depletion mutant strain CNV6 where each homolog of chromosome 7 is marked by the auxotrophic marker *HIS1* or *ARG4* (Fig. 6C). Natural rate of loss of a chromosome in the wild-type strain RM1000AH of *C. albicans* is <1 × 10^−3^/ cell/generation (Varshney *et al.*, 2015). We observed that Ipl1-depleted CNV6 mutant cells exhibited loss of chromosome 7 at a frequency of 8.2 × 10^−3^/cell/generation. Such a higher incidence of chromosome loss in the Ipl1 mutant cells explains the occurrence of aneuploid cells in the population resulting in FLC resistance. Recently, chromosome 7 trisomies have been linked to increased fitness of the pathogen in the human gut niche (Ene *et al.*, 2018). Taken together, these results indicate that Ipl1 is required for timely and proper chromosome segregation during the cell division to prevent aneuploidy-associated drug resistance in *C. albicans*.

**Figure 6.**
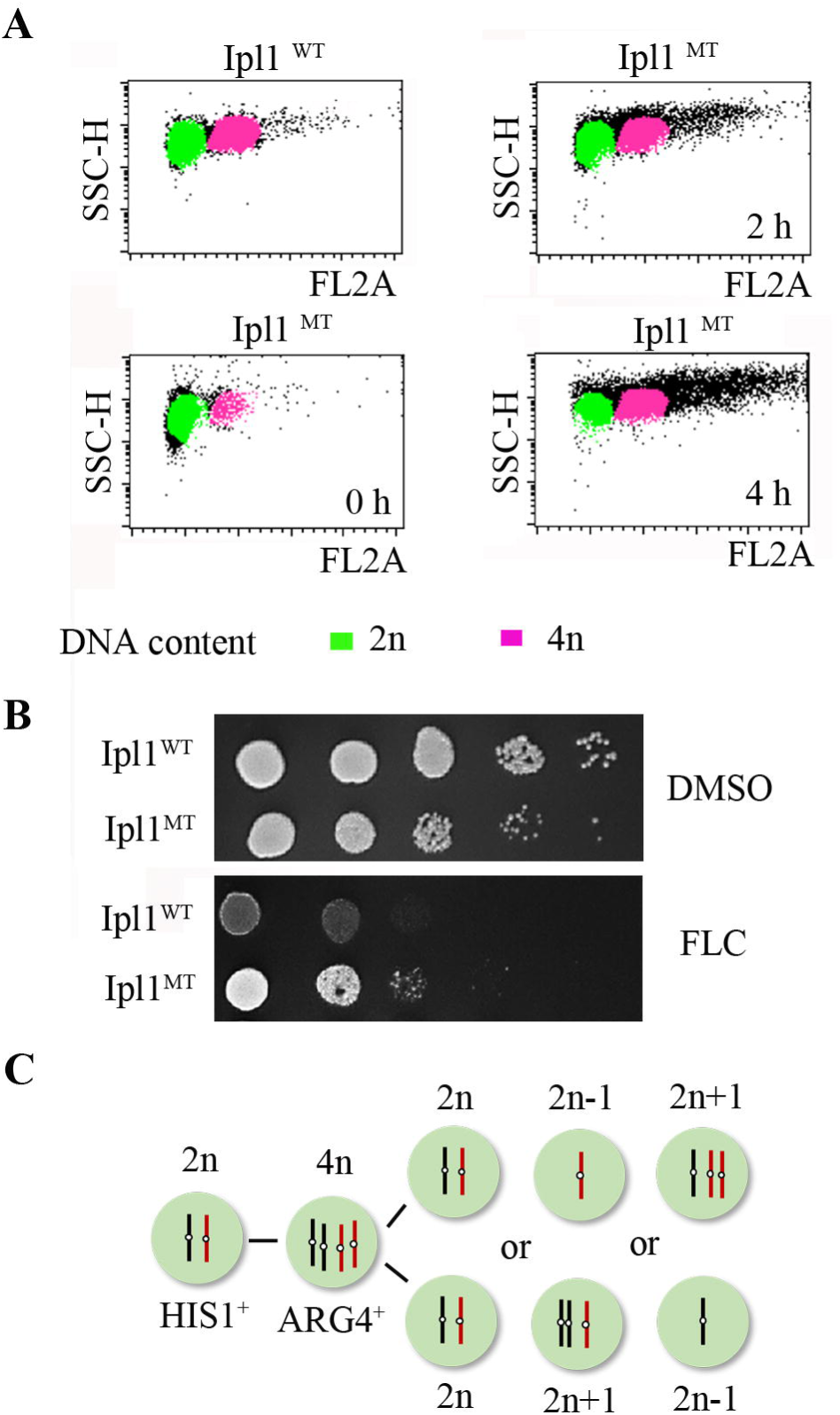
Altered ploidy in Ipl1-depleted cells results in drug resistance. **(A)** Dot-plots representing the altered morphology and DNA content measured in the wild-type and Ipl1-depleted cells using flow cytometry after growth in the non-permissive conditions at the indicated time-points. **(B)** Anti-fungal drug resistance assays corroborating aneuploidy in the Ipl1-depleted cells grown in non-permissive conditions for 4 h. The wild-type and Ipl1-depleted cells were 10-fold serially diluted and spotted on the non-permissive medium containing FLC (64 µg/ml). **(C)** Schematic depicting mis-segregation of chromosome 7.

## Discussion

In this work, we show that Aurora B kinase Ipl1 localizes to the kinetochores from the G1/S phase till metaphase and associates with the mitotic spindle from metaphase till the mitotic spindle disassembles in anaphase. Ipl1 possesses an unusual activation loop, is required for cellular viability and morphogenesis in polymorphic budding yeast *C. albicans*. Ipl1 plays a critical role in orchestrating the MT dynamics and facilitates the timely separation of SPBs. Even though Ipl1 is not directly involved in maintaining structural integrity and clustering of kinetochores in *C. albicans*, it is required for the maintenance of kinetochore geometry to form bilobed structures along the mitotic spindle to facilitate spindle bi-orientation. As expected Ipl1-depletion results in untimely nuclear division leading to erroneous kinetochore-MT attachments resulting in aneuploidy-associated anti-fungal drug resistance. Taken together, Ipl1 regulates the process of chromosome segregation by correcting erroneous kinetochore-MT attachments and modulating the synchronous movement of kinetochores along the mitotic spindle in the human pathogen *C. albicans* (Fig. 7).

**Figure 7.**
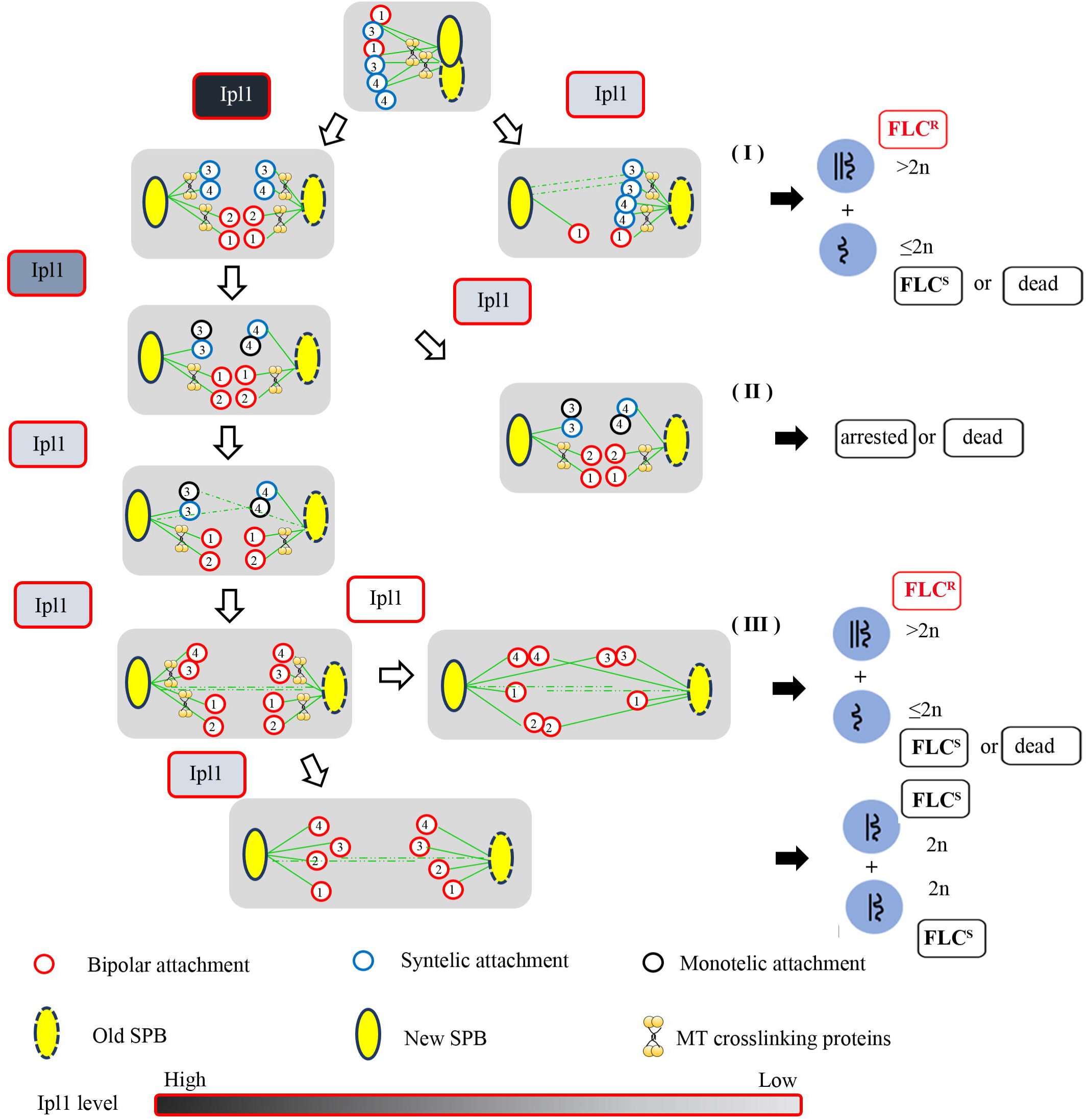
Model showing the role of Ipl1 in bipolarity establishment in *C. albicans*. Schematic of formation of equal bipolar kinetochore clusters during mitotic progression in the wild-type cells of *C. albicans* (above). The activity of Ipl1 (high, medium and low) is differentially regulated during different stage of the cell cycle, shown in the gradient colors of grey. Reduction in the activity of Ipl1 at different stages results in accumulation of cells with multiple defects, (I) accumulation of cells with syntelic attachments resulting in unequal distribution of kinetochore clusters, (II) accumulation of cells with monotelic attachments resulting in formation of monopolar kinetochore cluster and (III) accumulation of cells with asynchronous arrangement of kinetochores resulting disorganised kinetochores clusters.

MTs are highly dynamic structures that control cell shape, cell division, motility, and differentiation (Poulain & Sobel, 2010). The dynamics of MTs is modulated by MT regulatory proteins such as MAPs, +TIPs, and motor proteins. Several kinases spatiotemporally regulate the activities of these proteins for timely execution of the cell cycle. The spatiotemporal regulation by these kinases differs as some of the proteins are sequestered in different cellular compartments such as the nucleus or cytoplasm during various stages of cell cycle. (Janke, 2014, Wloga & Gaertig, 2010, Janke & Bulinski, 2011). In different organisms, activities of MT regulatory proteins have been shown to be tightly regulated by Aurora kinase for the proper execution of mitosis (Blangy *et al.*, 1995, Ohi *et al.*, 2004). Here, we show the role of Aurora kinase/Ipl1 in the spatiotemporal regulation of the MT dynamics in an ascomyceteous pathogenic polymorphic fungus *C. albicans*, which has one of the smallest known eukaryotic mitotic spindles of ~ 0.8 µm (McCoy *et al.*, 2015, Berman, 2006, Brand, 2012). It exhibits various cellular morphologies and provides an opportunity to determine the role of mitotic spindle properties in regulating the cell dimensions. It is an ideal system to explore various features of MT dynamics during mitosis at relatively smaller length scales where aneuploidy is tolerated under harsh host conditions.

While exploring the possible roles of Ipl1 in orchestrating MT dynamics during different stages of cell cycle, we observed that Ipl1 depleted cells have defective spindle morphologies. Further analysis of the SPB-to-SPB distance in the wild-type and Ipl1-depleted cells revealed that Ipl1 depletion leads to an aberrant SPB separation which may result in a mitotic delay or cell death. On the contrary, *ipl1-2* cells of *S. cerevisiae* do not exhibit any gross morphological differences in the spindle morphology (Biggins *et al.*, 1999). In addition, *ipl1-2* cells exhibit no difference in the separation of the SPBs throughout the cell cycle as compared to the wild-type (Biggins *et al.*, 1999). Interestingly, another allele of Ipl1, *ipl1-315* is required for the centrosome-mediated process of spindle assembly in the absence of the BimC motor protein, Cin8 (Kotwaliwale *et al.*, 2007). Aurora-B kinase also facilitates chromatin-mediated spindle assembly by inhibiting MCAK in vertebrates (Sampath *et al.*, 2004). Therefore, we speculate that in *C. albicans*, Ipl1 regulates the chromatin-mediated spindle assembly through phosphorylation of the yeast MCAK-like protein Kip3 or facilitates MTOC-mediated spindle assembly by phosphorylating kinesin-5 motors such as Cin8. It is noteworthy that the *C. albicans* genome has only one member of the kinesin-5 motor, i.e., Kip1 (Chua *et al.*, 2007) and an additional kinesin motor, Kip99. Kip1 deletion mutants in *C. albicans* exhibit polarised growth and defects in SPB separation (Chua *et al.*, 2007).

*ipl1-321* cells in *S. cerevisiae* display highly asymmetric distribution of Ndc80-GFP signals indicating presence of syntelic attachments in the cell (Marco *et al.*, 2013). Under the influence of Ipl1, syntelic attachments get corrected and form more stable bipolar attachments (Tanaka. *et al.*, 2002, Pinsky & Biggins, 2005, Pinsky *et al.*, 2006). Ipl1-depleted cells in *C. albicans* results in heterogeneity in the phenotype of kinetochore distribution suggesting functions of Ipl1 at different stages of cell cycle. We speculate that in addition to the well-conserved function of Ipl1 in correcting syntelic attachments, Ipl1 may also be involved in recapturing of monotelic attachments and maintenance of bilobed kinetochore organisation along the mitotic spindle in *C. albicans*.

Mutations in the essential gene *IPL1* results in aneuploidy and /or elevated ploidy in *S. cerevisiae* (Zhu *et al.*, 2016). Interestingly, acquisition of aneuploidy provides fitness to another ascomyceteous budding yeast *C. albicans* in response to anti-fungal drugs such as FLC (Gerstein & Berman, 2015). *C. albicans* cells acquire multiple aneuploid chromosomes just after few generations when grown in the presence of FLC (Selmecki *et al.*, 2009). We find that reduced levels of Ipl1 in *C. albicans* results in altered ploidy and exhibit FLC resistance. It is possible that the clinical strains isolated from the patients treated with anti-fungal drugs acquire mutations in *IPL1* which enable it to survive better in the host environment. Together, Ipl1 spatiotemporally modulates the dynamics of ipMTs/kMTs possibly by regulating kinesin-related motors and ensures bipolar spindle formation and bilobed geometry of kinetochores to prevent aneuploidy in the pathogenic budding yeast *C. albicans*.

## Experimental procedures

### Yeast strains, plasmids, and media conditions

Strains and primers used in this study are listed in the Supporting Information Table S1 and Supporting Information Table S2 respectively.

### Construction of the conditional mutant of IPL1 in RM1000AH

The conditional mutant strains of *IPL1* were constructed by deleting the first allele of *IPL1* (C6_02320C) with the Ca*NAT* flipper cassette (Reuss *et al.*, 2004) and by replacing the promoter of the second allele with the *PCK1* promoter (Leuker *et al.*, 1997). To delete the first copy, a Ca*NAT* flipper deletion cassette was constructed using the plasmid pSFS2a. The 3’ untranslated region (UTR) of the *IPL1* ORF was amplified from the SC5413 genomic DNA using oligos (NV13 and NV14) listed in the Supporting Information Table S2 and cloned into NotI and SacII sites of the plasmid pSFS2a to obtain pNV1. The 3’ UTR of the *IPL1* ORF was amplified using oligos NV11 and NV12 and cloned into KpnI and XhoI sites of pNV1 to obtain pNV2. The strain RM1000AH was transformed with pNV2 digested with KpnI and SacII by the lithium acetate procedure and the transformants were selected on YPDU plates containing nourseothricin (100 µg/ml). The transformants were screened for the stable integrants of digested linear DNA fragments into the genome at the right locus first by PCR using oligos (NV22 and NATmidF) and subsequently by Southern hybridization using a probe region amplified by NV76 and NV77 (Supporting Information Fig. S3A). The correct transformants (CNV4) were grown in YPM (1% yeast extract, 2% peptone, 2% maltose) supplemented with uridine (0.1 µg/ml) and plated on YPDU plates to recycle the *NAT* marker. Single colonies were replica plated on YPDU and YPDU + NAT (100 µg/ml) plates. Nou^s^ colonies were selected and confirmed by Southern hybridization for the first copy *ipl1* deletion followed by removal of the marker gene using a probe region (Supporting Information Table S2). In the resulting strains CNV5, the remaining wild-type allele of *IPL1* was placed under the control of the *PCK1* promoter. To generate the *PCK1*p*-IPL1* cassette, the 5’ UTR was amplified with primers NV63 and NV12 (Supporting Information Table S2) and cloned into KpnI and XhoI sites of pBS II KS (-) to obtain pNV7. The Ca*URA3* sequence was obtained as a HindIII digest from pCaURA3 and cloned into the HindIII site of pNV7 to obtain pNV8. Subsequently, the 5’ coding region of the *IPL1* ORF including the start codon was amplified with primers NV35 and NV36 (Supporting Information Table S2) and cloned into BamHI and NotI sites of pNV8. Finally, the *PCK1* promoter containing fragment obtained from pCAO1 (Leuker *et al.*, 1997) was cloned into the BamHI site of pNV9 to obtain the final construct as pNV10. pNV10 was digested with SacII and NotI and was used to transform CNV5 to give rise to CNV6 and CNV7 (see Supporting Information Table S1 for the genotypes of the strains). The conditional mutants were confirmed by PCR using oligos (NV88 and NV102) listed in the Table S2. The transformants were further confirmed by Southern hybridization using a probe region amplified by NV76 and NV77 (Supporting Information Fig. S3A).

### Construction of the conditional mutant of IPL1 in strains expressing GFP-tagged CENP-A, Dad2, and Tub4 co-expressing RFP-tagged Tub1 and protein A-tagged CENP-A

To study the dynamics of kinetochore proteins and SPBs, conditional *ipl1* mutants were constructed in the strains expressing GFP tagged CENP-A, Dad2, Tub1 and Tub4 respectively. To study the enrichment of CENP-A at the *CEN*, the conditional *ipl1* mutant was constructed in the strain containing protein-A-tagged CENP-A. The first copy of *IPL1* was replaced with a recyclable *NAT* in the strains, YJB8675, YJB10742, LSK111 and CAKS102 (see Supporting Information Table S1 for genotypes). In the resulting strains, CNV25, CNV18, CNV10, and CNV15 respectively, the *NAT* marker was first recycled and the remaining wild-type copy of *IPL1* was placed under the *PCK1* promoter. To construct the *PCK1*p*-IPL1* cassette with a *NAT* marker, the *PCK1*p*-IPL1* containing fragment was amplified from the genomic DNA of CNV5 using oligos (NV102 and NV103) listed in the Supporting Information Table S2 and cloned into the NotI and SacII sites of the plasmid pBSNAT to obtain pPCK1-IPL1-NAT. The plasmid was linearized with NsiI and was used to transform CNV26, CNV11, CNV19 and CNV16 to obtain CNV27, CNV12, CNV20 and CNV17, respectively. The strains were confirmed by PCR using oligos (NV88 and NV102) mentioned in the Supporting Information Table S2. Further, these conditional mutants were confirmed by their ability to form pseudo-hyphal cells under non-permissive media conditions.

To simultaneously localize tubulin in the strains, CNV14, CNV22, YJB10742 and LSK111, a plasmid containing 3’ UTR of *TUB1* cloned in frame with *RFP* harboring *HIS1* marker was linearized with XbaI. The linearized DNA was transformed into YJB10742, CNV12, LSK111, and CNV20 to obtain CNV21, CNV14, CNV13 and CNV22 respectively. The transformed were screened by microscopy for the correct integration of the cassette.

### Construction of the conditional mutant of IPL1 in strains expressing GFP-tagged Tub1

The conditional mutant of *IPL1* was constructed in the strain carrying GFP-tagged *Tub1* and RFP-tagged *Nop1*. To delete the first allele, an *IPL1* deletion cassette was constructed with *ARG4* as the selection marker. The recyclable *NAT* marker present in pNV2 was replaced by *ARG4* using oligos (NV251 and NV252) listed in the Supporting Information Table S2 to construct pNV2ARG4. The plasmid was digested with KpnI and SacII, and transformed into 12856 to obtain CNV8. The correct integrants were confirmed by PCR using oligos, NV22 and NV271 mentioned in the Supporting Information Table S2. The second allele of *IPL1* was placed under the *PCK1* promoter using *HIS1*. The *PCK1*p*-IPL1* fragment was amplified using primers, NV102 and NV103, from the genomic DNA of CNV5 and cloned into the NotI and SacII sites of pBSHIS to construct pBSPCK1IPL1HIS. The plasmid was linearized and used to transform CNV8 to obtain CNV9. The resulting strain was confirmed by their ability to form pseudo-hyphal cells under non-permissive conditions.

### Construction of a strain expressing double GFP epitope-tagged Ipl1 under the native promoter and Ndc80 tagged with RFP

To study the subcellular localization of Ipl1 under the native promoter, we constructed the strain expressing Ipl1-tagged with a double GFP at its C-terminus under the native promoter in the strain CNV2. The 3’coding sequence of *IPL1* excluding the stop codon was amplified with the oligos (NV124 and NV125) listed in the Supporting Information Table S2 and cloned into the NotI and SpeI sites of pBSGFPHIS1, to construct pBSIPL1GFPHIS1. Another fragment of *GFP* ORF was amplified using oligos (NV250 and SR67) and inserted into the SpeI site of pBSIPL1GFPHIS1 to obtain pBSIPL1-2GFP. After confirming the orientation of GFP by HpaI, the plasmid was linearized with XbaI and was used to transform to obtain CNV2. Further, to simultaneously localize IPL1 and Ndc80, we constructed a plasmid where the C-terminus of Ndc80 was cloned into SacII and SpeI sites of pRFPARG4 using oligos (NV448 and NV449) listed in the Table S2. The resulting plasmid pNdc80-RFP-ARG4 was linearized with XhoI and was used to transform the strain CNV2 to obtain CNV3. The transformants were screened for the presence of RFP tagged Ndc80 by microscopy.

### Construction of a strain expressing IPL1-TAP

To study the dynamic subcellular localization of Ipl1 and study its expression, we constructed the strain expressing *IPL1* tagged with a Protein-A epitope (Puig *et al.*, 2001) under its native promoter (CNV1). The 3’ UTR was amplified using oligos listed in the Supporting Information Table S2 (NV15 and NV16) and cloned into the SpeI and NotI sites of pBS II KS (-) to obtain pBSDSTAP. Subsequently, the Protein-A epitope with an auxotrophic marker Ca*URA3,* amplified from the plasmid pPK335 (Puig *et al.*, 2001) by oligos NV17 and NV18, listed in the Supporting Information Table S2 and 5’ UTR amplified from the SC5314 genomic DNA using oligos NV19 and NV20, listed in the Supporting Information Table S2 were cloned into BamHI and SpeI, and XhoI and BamHI sites of pBSDSTAP respectively, as a three-piece ligation reaction to obtain pBSIPL1TAPDS. The resulting plasmid was digested with XhoI and NotI and was used to transform BWP17 to obtain CNV1. The correct integrants were screened by PCR using oligos (NV34 and NV11) listed in the Supporting Information Table S2. The expression of Ipl1-Prot-A as a fusion protein was confirmed by western blotting using anti-protein A antibodies.

### Construction of the conditional mutant of IPL1 in a strain expressing GFP-tagged histone H2B

The conditional mutant of *IPL1* was constructed in the strain carrying GFP-tagged *H2B* (RSY15) obtained from Bennett’s Laboratory (Sherwood & Bennett, 2008). To delete the first allele, an *IPL1* deletion cassette constructed with *ARG4* as the selection marker, pNV2ARG4 (construction described previously) was used. The plasmid pNV2ARG4 was digested with KpnI and SacII, and transformed into strain RSY15 to obtain CNV23. The second allele of *IPL1* was placed under the *PCK1* promoter using PCK1p-HIS1 cassette. The plasmid pBSPCK1IPL1HIS was linearized and used to transform CNV23 to obtain CNV24. The conditional mutant strain was further confirmed by their ability to form pseudo-hyphal cells under non-permissive conditions.

### Media and growth conditions

The conditional mutant strains carrying *IPL1* under the control of the *PCK1* promoter were grown in YPS (1% yeast extract, 2% peptone, 2% succinate) as a permissive medium and YPD (1% yeast extract, 2% peptone, 2% dextrose) as a non-permissive medium. All the *C. albicans* strains were grown at 30°C.

### Live-cell imaging

The conditional mutant strains were grown overnight in the permissive medium and re-inoculated in the non-permissive media for 4 h. These cells were pelleted at 4,000 rpm and washed once with 1x phosphate buffered saline (PBS). The cell suspension was placed on the slide containing a thin 2% agarose patch prepared in dextrose and the patch was covered with a coverslip. Live-cell imaging was performed at 30°C on an inverted confocal microscope (ZEISS, LSM-880) equipped with a temperature-control chamber (Pecon incubator, XL multi SL), a Plan Apochromat 100x NA oil 1.4 objective and GaAsp photodetectors. For time-lapse microscopy of histone H2B-GFP, images were collected at a 60-s interval with 0.2% intensity exposure with 0.5 µm Z-steps using GFP filter lines (GFP/FITC 488 for excitation and GFP/FITC 500/550 band-pass for emission). All the images were displayed after the maximum intensity projection of images at each time using ImageJ.

### Microscopic image acquisition and processing

The conditional mutant strains were grown till OD_600_=1 in the permissive medium and re-inoculated in the non-permissive media for 8 h. These cells were pelleted at 4,000 rpm and washed once with 1x phosphate buffered saline (PBS) before the cell suspension was placed on a thin growth medium containing 2% agarose patch present on the slide. A coverslip was placed on the patch and processed for imaging. The images were acquired at room temperature using laser scanning inverted confocal microscope LSM 880-Airyscan (ZEISS, Plan Apochromat 63x, NA oil 1.4) or Leica SP8. The filters used were GFP/FITC 488, mCherry 561 for excitation. GFP/FITC 500/550 band pass or GFP 495/500 and mCherry 565/650 or mCherry 580-750 band pass for emission. Z-stack images were taken at every 0.3 μm and processed using ZEISS Zen software/ImageJ. All the images were digitally altered with minimal adjustments to levels and linear contrast till the signals were highlighted.

### Post-acquisition analysis

The distance between the two spindle pole bodies was measured after the 3D rendering of confocal images with IMARIS 7.6.4 software (Bitplane, Zurich, Switzerland). Images were filtered by Gaussian smoothing and a surface was created using a threshold of absolute intensity. Radii of the SPB spots were determined by taking a half of the longest diameters for each SPB spot measured in an individual stack in Imaris. The distances were obtained in statistics of processed images and these values were used to calculate SPB-to-SPB distance.

### Immunoblotting

Wild-type and mutant cells were grown under permissive and non-permissive conditions for the indicated time points. The cells of OD_600_=3 were harvested and washed with 1x PBS. The cells were resuspended in 12.5% TCA and lysates were precipitated at 13000 rpm for 10 min and washed with 80% acetone. The pellet was dried and resuspended in the lysis buffer (0.1 N NaOH, 1% SDS). The samples were diluted in 5x SDS loading dye (5% β-mercaptoethanol, 0.02% Bromophenol blue, 30% glycerol, 10% SDS, 250 mM Tris-Cl pH 6.8) and denatured. Denatured samples were subjected to electrophoresis using 10% SDS PAGE and transferred to a nitrocellulose membrane for 45 min at 25 V by semidry method (Bio-Rad). The membranes were blocked with 5% skim milk containing 1x PBS (pH 7.4) for 1 h at the room temperature followed by its incubation with primary antibodies in 2.5% skim milk overnight at 4°C. After three 10 min washes in PBST (1x PBS, 0.05% Tween) solution, the membranes were incubated in solutions containing secondary antibodies in 2.5% skim milk for 2 h. The membranes were washed with PBST (1X PBS, 0.05% Tween) solution thrice and the signals were detected using chemiluminescence method (SuperSignal West Pico Chemiluminescent Substrate, Thermo Scientific, Cat. No. 34080).

### Southern blotting

A high-quality genomic DNA was isolated from all the samples and quantified using the Quantity One software (Bio-Rad). An equal amount of DNA was digested with a suitable restriction enzyme and resolved on 1% agarose gel in 1x TAE buffer. The gel containing the resolved DNA was placed on a graduated scale and was imaged using Quantity One software (Bio-Rad). Subsequently, the gel was sequentially treated with 0.25 M HCl for acid nicking of DNA, denaturation buffer and, neutralization buffer. Acid nicked denatured DNA was then transferred to a Zeta-probe GT Nylon membrane in 10x SSC Buffer (Saline-Sodium Citrate) by capillary transfer method for 24-48 h. Following transfer, the membrane was washed with 2x SSC, dried and exposed to UV for 5 min in Genelinker (Bio-Rad) (12000 µJ x 100) for cross-linking DNA with the membrane. The blot was pre-hybridized for 2-4 h in 10 ml of 1x Southern hybridization buffer at 65°C. Meanwhile, a probe was radio-labeled with α ^32^ P dCTP using Klenow polymerase and random primer. The radio-labeled probe was prepared using a random primer labeling kit (BRIT-LCK-2). First, a mixture of 2 μl 50 ng PCR amplified DNA and 17 μl distilled water was boiled and chilled immediately for 5 min. Further, other reagents such as 5 μl random primer buffer, 12 μl of a cocktail of dNTPs except the radioactive dNTP, 5 μl α ^32^ P dCTP and 2 μl Klenow polymerase enzyme were added to the mixture and the reaction mixture was incubated at 37°C for 30 min. The reaction was stopped by adding 2 μl 0.5 M EDTA. After adding 50 μl 1x TE, the labeled probe was purified by passing through a Sephadex-G-50 column. The pre-hybridized blot was incubated with the purified radio-labeled probe overnight at 65°C. The blot was rinsed with wash buffer I once followed by wash buffer II, thrice at 65°C. The hybridized membrane was exposed to Phosphorimager films and images captured by Phosphorimager using the Image Reader FL2A5000 Ver.2 software.

### Flow cytometry

A 5 ml culture of *C. albicans* cells grown till OD_600_=0.5 was harvested, washed, resuspended in 100 μl sterilized water and fixed with 1 ml 70% ethanol overnight at 4°C. Cells were washed and resuspended in 100 μl of RNase buffer. Cells were then treated with 10 μl RNase A (10 mg/ml) for 4 h at 37°C. Cells were washed with 1x PBS resuspended in 900 μl of 1x PBS and incubated overnight at 4°C. Cells were stained with (0.005 μg/ml) propidium iodide for 30 min in dark conditions, just prior to the data acquisition. After a brief sonication (10 amp for 10 sec), data from 10,000 cells per time point were collected using a FACS Calibur.

### Chromatin-immunoprecipitation (ChIP)

Chromatin-immunoprecipitation (ChIP) followed by PCR was done as mentioned previously (Sanyal *et al.*, 2004). An asynchronous culture was grown in YPD till OD600 =1.0. Cells were cross-linked with 37% formaldehyde for 15 min. Next, the samples were quenched by adding 125 mM glycine. Further, the sonication was performed with Biorupter (Diagenode) to obtain sheared chromatin fragments of an average size of 300-500 bp. The fragments were immuno-precipitated with anti-Protein-A antibodies at a working concentration of 20 µg/ml. The immune-precipitated fragments were next incubated with the Protein-A sepharose beads. The ChIP DNA obtained from anti-protein-A ChIP assays were analyzed by qPCR and oligos that amplify central regions of *CEN1* and *CEN7*. The presence of background immuno-precipitated DNA was detected by qPCR and oligos that amplify non-centromeric DNA. For anti-Protein-A ChIP analysis, qPCR was performed on a Rotor-Gene 6000 real-time PCR machine with iQ^TM^ SYBR Green Super Mix. The cycling parameters used are as follows: 94°C for 30 s, 55°C for 30 s and 72°C for 45 s repeated 40 times. The CENP-A enrichment was quantified by percent input method.

### Antibodies

Primary antibodies used for western blot analysis were mouse anti-protein A antibodies (dilution 1:5000) (Sigma, Cat. No. P3775) and mouse anti-PSTAIRE (dilution 1:2000) (Abcam, Cat. No.10345). Secondary antibodies used are goat anti-mouse HRP conjugated antibodies (dilution 1:10,000) (Bangalore Genei, Cat. No. HP06).

### Anti-fungal drug resistance assay

The wild-type and Ipl1-depleted cells grown in the non-permissive condition for 4 h were 10-fold serially diluted and spotted on plates containing the non-permissive medium having indicated concentrations of fluconazole (Sigma, Cat. No. F8929-100 MG) and incubated further at 30°C for the antifungal drug resistance assay.

### Statistical analyses

Statistical differences were determined using paired Student’s *t*-test to calculate the statistical significance using a column type mean with the SD or SEM using Prism 7 software.

## Acknowledgments

We thank B. Suma and S. Kumar at the confocal imaging facility of the Molecular Biology and Genetics Unit, JNCASR, and P. Iyer at the imaging facility at ACTREC, Navi Mumbai, India for assisting in image analysis. We thank Aswathy Narayanan for performing spotting experiments. We thank J Berman, Tel Aviv University and R. Bennett, Brown University for providing us with strains and J. Morschhӓuser, Universität Würzburg for the *SAT1* flipper cassette plasmid. This work was partially supported by the Tata Innovation Fellowship (BT/HRD/35/01/03/2017) and intramural financial support from JNCASR to KS. NV was supported by a fellowship from the Council of Scientific & Industrial Research (CSIR), India. The authors declare no competing interests.

## Authors contributions

Conceived and designed the experiments: NV and KS. Contributed reagents and conducted the experiments: NV. Analyzed the data: NV and KS. Wrote the paper: NV and KS.

