## Supplemental file for "Aurora kinase Ipl1 facilitates bilobed distribution of clustered kinetochores to ensure error-free chromosome segregation in *Candida albicans*"

### Supporting Figures

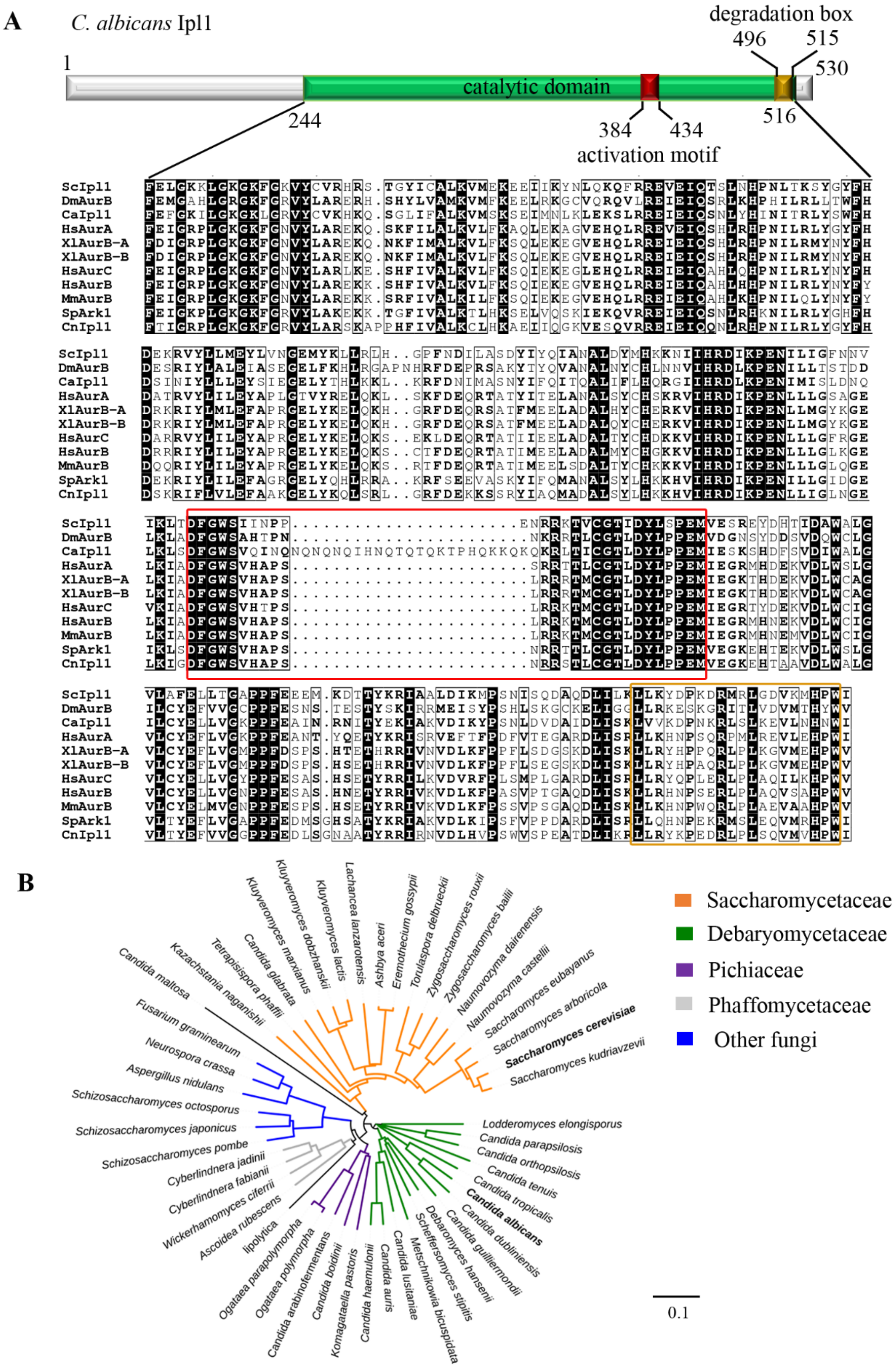

**Figure S1. Sequence conservation and domain analysis of Aurora kinases. (A)** Domain analysis of Ipl1 protein sequence in *C. albicans*. The catalytic domain (amino acids 244-516) is shown in green having a unique activation motif (amino acids 384-434) shown in red and degradation box (amino acids 496-515) shown in yellow. Comparison of the catalytic domain of the Aurora kinase B homolog Ipl1 in *C. albicans* with the catalytic domains of the Aurora/Ipl1 protein-related kinases from different species. The conserved residues are shaded in black and dashes are the gaps introduced to maximize alignment. CaIpl1 contains an additional stretch of amino acids within the activation motif (DFGSXXXXXXXXRXTXCGTXDYLPPE) indicated in the red box which is a conserved signature for Aurora/Ipl1 family members. The catalytic domain also contains a conserved degradation box (D box) (LLXXXPXXRXXLXXXXXHPW) near its C-terminus (yellow box). Sc: *Saccharomyces cerevisiae*, Dm: *Drosophila melanogaster*, Ca: *Candida albicans*, Hs: *Homo sapiens*, Xl: *Xenopus laevis*, Mm: *Mus musculus*, Sp: *Schizosaccharomyces pombe*, Cn: *Cryptococcus neoformans*. (**B)** Phylogenetic relationship of catalytic domain of Ipl1 from Saccharomycotina yeasts inferred from the alignment data obtained through Neighbourhood joining method using Simple Phylogeny and iTOL (Interactive Tree Of Life). Amino acid sequences of Ipl1 from different species were retrieved from NCBI-BLASTp using *S. cerevisiae* Ipl1 as a query and the catalytic domain of Ipl1 was aligned using Clustal Omega. The position of aurora kinase homolog in *C. albicans and S. cerevisiae* is shown in bold.

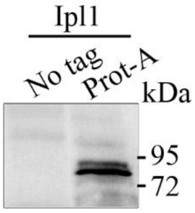


**Figure S2. Expression analysis of Protein A-tagged Ipl1 by western blotting.** The Protein A-tagged Ipl1 was pulled down using IgG sepharose 6 fast flow beads from a 500 ml culture of $OD_{600}$=1.0. The expression of Protein A-tagged Ipl1 was verified by western blotting using anti-Protein A antibodies. Protein A-tagged Ipl1 runs at a position between 95 kDa and 72 kDa molecular weight (MW) in a denaturing SDS-PAGE. Parental strain BWP17 was used as an no tag control.

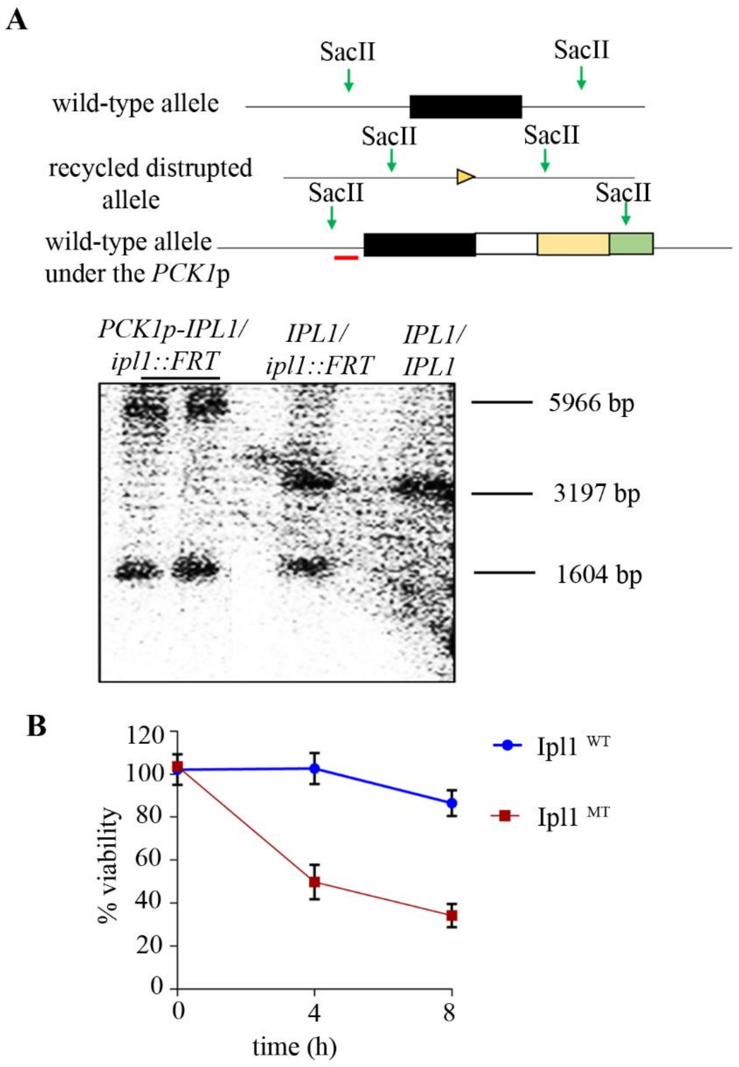


**Figure S3. Ipl1 is essential for the viability, depletion of which induces aberrant morphological states in *C. albicans*.** (**A)** Line diagrams showing the Southern hybridization strategy to confirm the construction of the conditional mutant of *IPL1* with the *PCK1* promoter, indicating the SacII restriction enzyme sites with green arrows and the position of the probe used for hybridization with a red bar. The phosphorimager image shows Southern hybridization results of SacII digested DNA from the wild-type strain (*IPL1/IPL1*), *NAT*$^{S}$ heterozygous deletion mutant strain (*IPL1/ipl1::FRT*) and two independent conditional *IPL1* mutant strains (*PCK1*p-*IPL1/ipl1::FRT*). (**B)** A graph depicting a drop-in the viability of cells in the Ipl1 mutant after their growth in non-permissive conditions for 4 h and 8 h as compared to the wild-type.

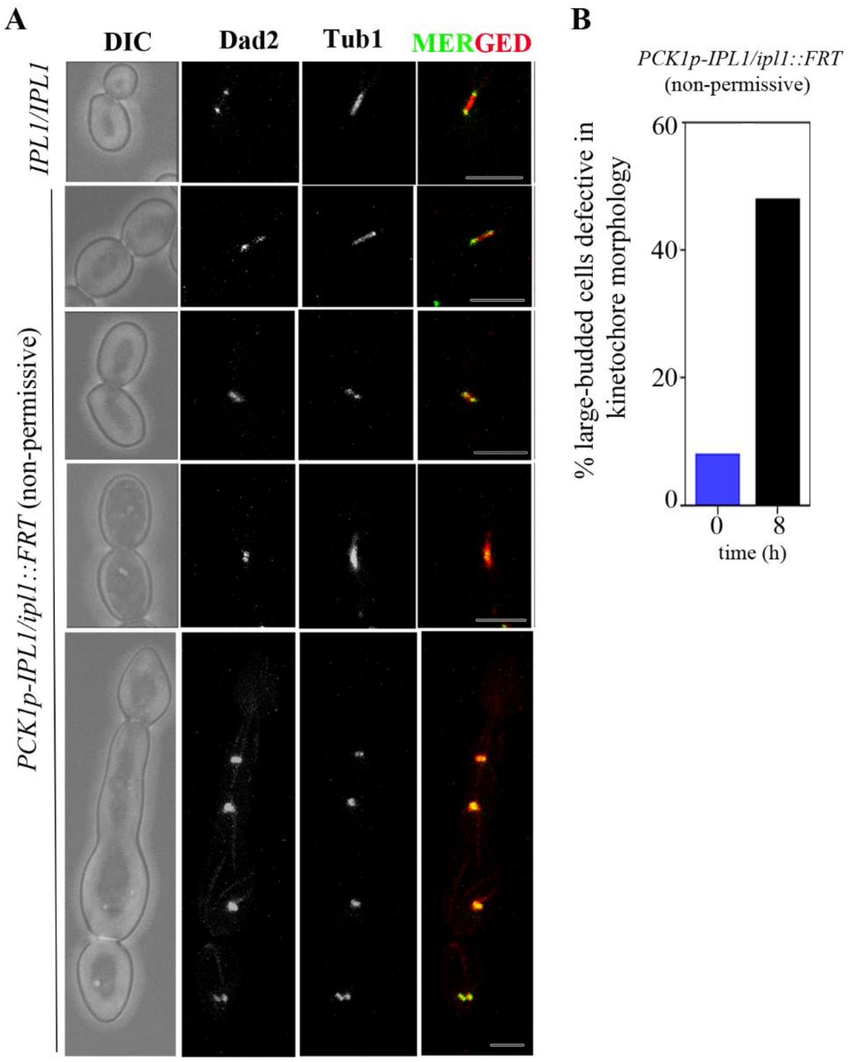


**Figure S4. Disorganization of the kinetochores and improper kinetochore-micrptubule attachments in the Ipl1-depleted cells. (A)** Images of the organization of kinetochores (Dad2-GFP) along the spindle axis (RFP-Tub1) in the wild-type CNV21 and mutant CNV22 cells of *IPL1* after growth in the non-permissive conditions for 8 h. Bar, 5 µm. (**B)** Quantification of large-budded cells in the Ipl1 mutant having disorganized kinetochores or improper kinetochore-MT attachments after growth in the non-permissive conditions for 8 h.

### Supporting tables

**Table S1. Strains used in this study**

| Name | Parent | Genotype | Reference |
|---|---|---|---|
| **RM1000AH** | RM1000 | *Δura3::imm434/Δura3::imm434Δhis1::hisG/ Δhis1::hisG arg4::HIS1/ARG4* | (Sanyal *et al.*, 2004) |
| **BWP17** | RM1000 | Δ*ura3::*imm434/Δ*ura3::*imm434 Δ*his1::*hisG/ Δ*his1::*hisG Δ*arg4::*hisG/ Δ*arg4::*hisG | (Wilson *et al.*, 1999) |
| **SN148** | | Δ*ura3::*imm434/Δ*ura3::*imm434 Δ*his1::*hisG/ Δ*his1::*hisG Δ*arg4::*hisG/ Δ*arg4::*hisG Δleu2*::*hisG/ Δleu2*::*hisG | (Noble & Johnson, 2005) |
| **CNV1** | BWP17 | Δ*ura3::*imm434/Δ*ura3::*imm434 Δ*his1::*hisG/ Δ*his1::*hisG Δ*arg4::*hisG/ Δ*arg4::*hisG *IPL1-TAP-URA3/IPL1* | This study |
| **CNV2** | CNV2 | Δ*ura3::*imm434/Δ*ura3::*imm434 Δ*his1::*hisG/ Δ*his1::*hisG Δ*arg4::*hisG/ Δ*arg4::*hisG Δleu2*::*hisG/ Δleu2*::*hisG *ipl1:FRT/IPL1-2xGFP-HIS1* | This study |
| **CNV3** | CNV2 | Δ*ura3::*imm434/Δ*ura3::*imm434 Δ*his1::*hisG/ Δ*his1::*hisG Δ*arg4::*hisG/ Δ*arg4::*hisG Δleu2*::*hisG/ Δleu2*::*hisG *ipl1:FRT/IPL1-2xGFP-HIS1 NDC80/NDC80-RFP-ARG4* | This study |
| **CNV4** | RM1000AH | *Δura3::imm434/Δura3::imm434 Δhis1::hisG/ Δhis1::hisG arg4::HIS1/ARG4 ipl1::NAT-Flp/IPL1* | This study |
| **CNV5** | CNV4 | *Δura3::imm434/Δura3::imm434 Δhis1::hisG/ Δhis1::hisG arg4::HIS1/ARG4 ipl1::FRT/IPL1* | This study |
| **CNV6** | CNV5 | *Δura3::imm434/Δura3::imm434 Δhis1::hisG/ Δhis1::hisG arg4::HIS1/ARG4 ipl1::FRT/PCK1*p-*IPL1* | This study |
| **CNV7** | CNV6 | *Δura3::imm434/Δura3::imm434 Δhis1::hisG/ Δhis1::hisG arg4::HIS1/ARG4 ipl1::FRT/PCK1*p-*IPL1* | This study |
| **12865** | BWP17 | *Δura3::imm434/Δura3::imm434 Δhis1::hisG/ Δhis1::hisG Δarg4::hisG/ Δarg4::hisG TUB1-GFP URA3/TUB1 NOP1-RFP NAT/NOP1* | (Harrison *et al.*, 2014) |
| **CNV8** | 12865 | *Δura3::imm434/Δura3::imm434 Δhis1::hisG/ Δhis1::hisG Δarg4::hisG/ Δarg4::hisG TUB1-GFP URA3/TUB1 NOP1-RFP NAT/NOP1 ipl1::ARG4/IPL1* | This study |
| **CNV9** | CNV8 | *Δura3::imm434/Δura3::imm434 Δhis1::hisG/ Δhis1::hisG Δarg4::hisG/ Δarg4::hisG TUB1-GFP URA3/TUB1 NOP1-RFP NAT/NOP1 ipl1::ARG4/PCK1*p-*IPL1-HIS1* | This study |
| **LSK111** | SN148 | *Δura3::imm434/Δura3::imm434 Δhis1::hisG/ Δhis1::hisG Δarg4::hisG/ Δarg4::hisG Δleu2::hisG/ Δleu2::hisG TUB4-GFP-URA3/TUB4* | (Sutradhar *et al.*, 2015) |
| **CNV10** | LSK111 | *TUB4-GFP-URA3/TUB4 ipl1::NAT-Flp/IPL1* | This study |

| | | | |
|---|---|---|---|
| **CNV11** | CNV10 | *TUB4-GFP-URA3/TUB4 ipl1::FRT/IPL1* | This study |
| **CNV12** | CNV11 | *TUB4-GFP-URA3/TUB4 ipl1::FRT/PCK1p-IPL1-NAT* | This study |
| **CNV13** | LSK111 | *TUB4-GFP-URA3/TUB4 TUB1-mCherry-HIS1/TUB1* | This study |
| **CNV14** | CNV12 | *TUB4-GFP-URA3/TUB4 TUB1-mCherry-HIS1/TUB1 ipl1::FRT/PCK1*p-*IPL1-NAT* | This study |
| **CAKS102** | SN148 | *Δura3::imm434/Δura3::imm434,Δhis1::hisG/Δhis1::hisG, Δarg4::hisG/Δarg4::hisG, Δleu2::hisG/Δleu2::hisG CENP-A/CENP-A-TAP(URA3)* | (Mitra *et al.*, 2014) |
| **CNV15** | CAKS102 | *CENP-A/CENP-A-TAP(URA3) ipl1::NAT-Flp/IPL1* | This study |
| **CNV16** | CNV15 | *CENP-A/CENP-A-TAP(URA3) ipl1::FRT/IPL1* | This study |
| **CNV17** | CNV16 | *CENP-A/CENP-A-TAP(URA3) ipl1::FRT/PCK1p-IPL1-NAT* | This study |
| **YJB10742** | BWP17 | Δ*ura3::*imm434/Δ*ura3::*imm434 Δ*his1::*hisG/ Δ*his1::*hisG Δ*arg4::*hisG/ Δ*arg4::*hisG *DAD2-GFP-URA3/DAD2* | (Burrack *et al.*, 2011) |
| **CNV18** | YJB10742 | *DAD2-GFP-URA3/DAD2 ipl1::NAT-Flp/IPL1* | This study |
| **CNV19** | CNV22 | *DAD2-GFP-URA3/DAD2 ipl1::FRT/IPL1* | This study |
| **CNV20** | CNV23 | *DAD2-GFP-URA3/DAD2 ipl1::FRT/PCK1p-IPL1-NAT* | This study |
| **CNV21** | YJB10742 | Δ*ura3::*imm434/Δ*ura3::*imm434 Δ*his1::*hisG/ Δ*his1::*hisG Δ*arg4::*hisG/ Δ*arg4::*hisG *DAD2-GFP-URA3/DAD2 TUB1-mCherry-HIS1/TUB1* | This study |
| **CNV22** | CNV20 | Δ*ura3::*imm434/Δ*ura3::*imm434 Δ*his1::*hisG/ Δ*his1::*hisG Δ*arg4::*hisG/ Δ*arg4::*hisG *DAD2-GFP-URA3/DAD2 TUB1-mCherry-HIS1/TUB1 ipl1::FRT/PCK1*p-*IPL1-NAT* | This study |
| **RSY15** | RBY1132 | *leu2/leu2 his1/his1 arg4/arg4 HTB/HTB-GFP::SAT1* | (Sherwood & Bennett, 2008) |
| **CNV23** | RSY15 | *leu2/leu2 his1/his1 arg4/arg4 HTB/HTB-GFP::SAT1 ipl1::ARG4/IPL1* | This study |

| | | | |
|---|---|---|---|
| **CNV24** | CNV23 | *leu2*/*leu2 his1*/*his1 arg4*/*arg4 HTB*/*HTB-GFP*::*SAT1 ipl1::ARG4/PCK1p-IPL1-HIS1* | This study |
| **YJB8675** | BWP17 | *Δura3*::*imm434/Δura3*::*imm434, Δhis1*::*hisG/Δhis1*::*hisG Δarg4*::*hisG/Δarg4*::*hisG, CSE4/CSE4*-GFP-*CSE4* | (Joglekar *et al.*, 2008) |
| **CNV25** | CNV24 | *CSE4/CSE4*-GFP-*CSE4 ipl1::NAT-Flp/IPL1* | This study |
| **CNV26** | CNV25 | *CSE4/CSE4*-GFP-*CSE4 ipl1::FRT/IPL1* | This study |
| **CNV27** | CNV26 | *CSE4/CSE4*-GFP-*CSE4 ipl1::FRT/IPL1 ipl1::FRT/PCK1*p-*IPL1-NAT* | This study |

**Table S2. Primers used in this study**

| Primer name | Sequence | Description |
|---|---|---|
| NV11 | AGTGGTACCGAAACTACGCTAACCAAACTC | Forward cloning primer for upstream of *ipl1* deletion cassette |
| NV12 | CAGCTCGAGAATTGAGGACAGGCAGGT | Reverse cloning primer for upstream of *ipl1* deletion cassette and reverse cloning primer for upstream of *PCK1p-IPL1* cassette |
| NV13 | ATTAGCGGCCGCACGAAGGCAAATCACATCT | Forward cloning primer for downstream of *ipl1* deletion cassette |
| NV14 | TCCCCGCGGTCAGGGTCAATTGGTGGA | Reverse cloning primer for downstream of *ipl1* deletion cassette |
| NV15 | CAGCTCGAGCAACGTGGAATAATTCATCGTG | Forward cloning primer for downstream of *IPL1-TAP* cassette |
| NV16 | CGCGGATCCTTTGTAAATATTTTTTGGCCATTTTGG | Reverse cloning primer for downstream of *IPL1-TAP* cassette |
| NV17 | CGCGCAATTAACCCTCACTA | Forward cloning primer for *TAP-URA3* of *IPL1-TAP* cassette |
| NV18 | CGGACTAGTACGACTCACTATAGGGCGAA | Reverse cloning primer for *TAP-URA3* of *IPL1-TAP* cassette |
| NV19 | CGGACTAGTACGAAGGCAAATCACATCT | Forward cloning primer for upstream of *IPL1-TAP* cassette |
| NV20 | ATTAGCGGCCGCTCAGGGTCAATTGGTGGA | Reverse cloning primer for upstream of *IPL1-TAP* cassette |
| NATmidF | TTAGAGACACAAACGAACAATGTACC | Forward primer for confirmation of integration of *ipl1* deletion cassette |
| NV22 | ATTAGCGGCCGCTCAGGGTCAATTGGTGGA | Reverse primer for confirmation of integration of *ipl1* deletion cassette |
| NV34 | GAGCACGTATTGGGTTTGC | Reverse primer for confirmation of integration of *IPL1-TAP* cassette |
| NV35 | CGCGGATCCATGATGCTTCCACGTAACTCACC | Forward cloning primer for downstream of *PCK1*p-*IPL1* cassette |
| NV36 | ATTAGCGGCCGCGGGTAGCTGTTGTTGTGGTTCT | Reverse cloning primer for downstream of *PCK1*p-*IPL1* cassette |
| NV63 | AGTGGTACCCATTTGAGCGCTACCAAGAG | Reverse cloning primer of upstream of *PCK1p-IPL1* cassette |

| | | |
|---|---|---|
| NV76 | TTGTCGCCATAACATTTGTTG | Forward primer for probe used for confirmation of integration of *PCK1*p-*IPL1* |
| NV77 | CATAGATTTCTTCGAGCTCGT | Reverse primer for probe used for confirmation of integration of *PCK1*p-*IPL1* |
| NV88 | GCGTCGACTCATTTGTAAATATTTTTTG GCCATT | Reverse primer for confirmation of integration of *PCK1*p-*IPL1-NAT* cassette |
| NV102 | ATTAGCGGCCGCGGATCCACTGTATTCC AATTTA | Forward primer for confirmation of integration of *PCK1*p-*IPL1-NAT* cassette, Forward cloning primer for *PCK1*p-*IPL1 in* pBSNAT plasmid |
| NV103 | TCCCCGCGGGAATTTCAATTTCTCGTCG TAAAA | Reverse cloning primer for *PCK1*p-*IPL1 in* pBSNAT plasmid |
| NV251 | CAGCTCGAGCTTAACGCTTCAGGTTCCT G | Forward cloning primer for *ARG4* in *ipl1* deletion cassette |
| NV252 | ATTAGCGGCCGCGCAAATCTAAGAGTA GAGGCTA | Reverse cloning primer for *ARG4* in *ipl1* deletion cassette |
| NV271 | AACTAAATAAAAGCTTCTTCCATCA | Forward primer for confirmation of integration of *ipl1* deletion cassette with *ARG4* |
| p2488-1 | CACTCTGACCAAATTCTCGTTTCC | Forward primer for ChIP PCR at the *CEN1* |
| p2488-2 | GCAACATCCGAGTAAGGTTTGTGG | Reverse primer for ChIP PCR at the *CEN1* |
| nCEN7-3 | GCATACCTGACACTGTCGTT | Forward primer for ChIP PCR at the *CEN7* |
| nCEN7-4 | AACGGTGCTACGTTTTTTTA | Reverse primer for ChIP PCR at the *CEN7* |
| Non-CEN7a | ACTCGCCTTCCCCTCCTTTAAATAG | Forward primer for ChIP PCR at the non-*CEN7* |
| Non-CEN7b | CCACTACTACGACTGTGGATTCACT | Reverse primer for ChIP PCR at the non-*CEN7* |
| NV124 | ATTAGCGGCCGCGCATTGATTTTTTTAC ATCAACGT | Forward cloning primer for Ipl1-2GFP cassette |
| NV125 | CGGACTAGTTTTGTAAATATTTTTTGGC CATTTTGG | Reverse cloning primer for Ipl1-2GFP cassette |

| | | |
|---|---|---|
| NV250 | CGCACTAGTTTTGTATAGTTCATCCATGCC | Reverse cloning primer for GFP in Ipl1-2GFP cassette |
| NV448 | TCCCCGCGGGGATTTACAACAACGAGGTATATC | Forward cloning primer for Ndc80-RFP cassette |
| NV449 | CGGACTAGTCTCTAGCTGTTGTGATGCTTGTT | Reverse cloning primer for Ndc80-RFP cassette |
| SR67 | CGCACTAGTATGAGTAAGGGAGAAGAACTTTTCAC | Forward cloning primer for second GFP in Ipl1-GFP cassette |
| NV250 | CGCACTAGT TTTGTATAGTTCATCCATGCC | Reverse cloning primer for second GFP in Ipl1-GFP cassette |

#### Supporting movie legends

**S1 Movie. Nuclear segregation in the wild-type and Ipl1-depleted cells.** The nucleus is visualized by GFP-tagged histone H2B. Images were acquired every 60 s with confocal microscopy.
